# Spike glycoprotein and host cell determinants of SARS-CoV-2 entry and cytopathic effects

**DOI:** 10.1101/2020.10.22.351569

**Authors:** Hanh T. Nguyen, Shijian Zhang, Qian Wang, Saumya Anang, Jia Wang, Haitao Ding, John C. Kappes, Joseph Sodroski

**Author notes:** Corresponding author: Joseph G. Sodroski, M.D., Dana-Farber Cancer Institute, 450 Brookline Avenue, CLS 1010, Boston, MA 02215. Hanh T. Nguyen and Shijian Zhang contributed equally to this article. Author order was determined by mutual agreement.

## Abstract

SARS-CoV-2, a betacoronavirus, is the cause of the COVID-19 pandemic. The SARS-CoV-2 spike (S) glycoprotein trimer mediates virus entry into host cells and cytopathic effects. We studied the contribution of several S glycoprotein features to these functions, focusing on those that differ among related coronaviruses. Acquisition of the furin cleavage site by the SARS-CoV-2 S glycoprotein decreased virus stability and infectivity, but greatly enhanced the ability to form lethal syncytia. Notably, the D614G change found in globally predominant SARS-CoV-2 strains restored infectivity, modestly enhanced responsiveness to the ACE2 receptor and susceptibility to neutralizing sera, and tightened association of the S1 subunit with the trimer. Apparently, two unique features of the SARS-CoV-2 S glycoprotein, the furin cleavage site and D614G, have evolved to balance virus infectivity, stability, cytopathicity and antibody vulnerability. Although the endodomain (cytoplasmic tail) of the S2 subunit was not absolutely required for virus entry or syncytium formation, alteration of palmitoylated cysteine residues in the cytoplasmic tail decreased the efficiency of these processes. As proteolytic cleavage contributes to the activation of the SARS-CoV-2 S glycoprotein, we evaluated the ability of protease inhibitors to suppress S glycoprotein function. Matrix metalloprotease inhibitors suppressed S-mediated cell-cell fusion, but not virus entry. Synergy between inhibitors of matrix metalloproteases and TMPRSS2 suggests that both proteases can activate the S glycoprotein during the process of syncytium formation. These results provide insights into SARS-CoV-2 S glycoprotein-host cell interactions that likely contribute to the transmission and pathogenicity of this pandemic agent.

**IMPORTANCE:** The development of an effective and durable SARS-CoV-2 vaccine is essential for combating the growing COVID-19 pandemic. The SARS-CoV-2 spike (S) glycoprotein is the main target of neutralizing antibodies elicited during virus infection or following vaccination. Knowledge of the spike glycoprotein evolution, function and interactions with host factors will help researchers to develop effective vaccine immunogens and treatments. Here we identify key features of the spike glycoprotein, including the furin cleavage site and the D614G natural mutation, that modulate viral cytopathic effects, infectivity and sensitivity to inhibition. We also identify two inhibitors of host metalloproteases that block S-mediated cell-cell fusion, which contributes to the destruction of the virus-infected cell.

## INTRODUCTION

Coronaviruses are enveloped, positive-stranded RNA viruses that cause respiratory and digestive tract infections in animals and humans (1–4). Two betacoronaviruses, severe acute respiratory syndrome coronavirus (SARS-CoV-1) and Middle Eastern respiratory syndrome coronavirus (MERS-CoV), caused deadly but self-limited outbreaks of severe pneumonia and bronchiolitis in humans in 2002 and 2013, respectively (2,5). In late 2019, an emergent betacoronavirus, SARS-CoV-2, was shown to cause COVID-19, a severe respiratory disease in humans with 3-4% mortality (6–12). MERS-CoV, SARS-CoV-1 and SARS-CoV-2 are thought to have originated in bats, and infected humans either directly or through intermediate animal hosts (1,4,9,13–15). The efficient transmission of SARS-CoV-2 in humans has resulted in a pandemic that has led to over a million deaths and threatened global health, economies and quality of life (11,12,16). An immediate goal for the current SARS-CoV-2 pandemic is to identify treatments, including passively administered neutralizing antibodies, that could ameliorate COVID-19 disease and improve survival (17,18). Another urgent priority is the development of a vaccine that could protect against SARS-CoV-2 infection (19–21).

The SARS-CoV-2 spike glycoprotein (S gp) is the major target of virus-neutralizing antibodies that are thought to be important for vaccine-induced protection (17–32). The SARS-CoV-2 S gp mediates the entry of the virus into host cells and influences tissue tropism and pathogenesis (22,23,32–35). The trimeric S gp is a Class I fusion protein that is cleaved into the S1 and S2 glycoproteins, which associate non-covalently in the trimeric spike. The receptor-binding domain (RBD) (residues 331-528) of the S1 subunit binds the receptor, angiotensin-converting enzyme 2 (ACE2) (22,23,32,35–38). The S2 subunit, which contains a fusion peptide and two heptad repeat regions (HR1 and HR2), mediates fusion of the viral and target cell membranes (22,35,39,40). Following receptor binding of S1, cleavage of S2 at the S2’ site by host proteases (Cathepsin B/L, TMPRSS2) is thought to activate extensive and irreversible conformational changes in S2 required for membrane fusion (38,41,42). The interaction of the HR1 and HR2 regions of S2 results in the formation of a stable six-helix bundle that brings the viral and cell membranes into proximity, promoting virus entry (39,40).

S gp diversity contributes to host and tissue tropism, transmissibility and pathogenicity of coronaviruses (1–4,34). The S gp of SARS-CoV-2 is 79.6% identical in amino acid sequence to that of SARS-CoV-1 (8,9,15), but possesses notable unique features. First, during its evolution from an ancestral bat coronavirus, the SARS-CoV-2 S gp acquired a multibasic sequence at the S1/S2 junction that is suitable for cleavage by furin-like proteases (9,15,41,42). A similar cleavage site sequence is present in the MERS-CoV S gp but not in the SARS-CoV-1 S gp. Second, compared with the original source virus in Wuhan, China, the more prevalent SARS-CoV-2 variants emerging in the global pandemic substitute a glycine residue for aspartic acid 614 (D614G) in the S1 C-terminal domain (CTD2); the D614G change is associated with higher levels of virus replication in cultured cells (43,44). Third, although the cysteine-rich S2 endodomains of SARS-CoV-1 and SARS-CoV-2 are 97% identical, the latter has an additional cysteine residue in its cytoplasmic tail (15). The S2 endodomain of SARS-CoV-1 is palmitoylated and has been shown to contribute to S gp function and localization to virus assembly sites in the endoplasmic reticulum-Golgi intermediate complex (ERGIC) (45–50).

Here, we evaluate the phenotypes of SARS-CoV-2 S gp mutants with alterations in the unique features described above, as well as other features that are potentially important for function. These include: a) the two proposed proteolytic cleavage sites at the S1/S2 junction, and the S2′ cleavage site immediately N-terminal to the fusion peptide; b) the polymorphic residue Asp/Gly 614; c) the putative fusion peptide in the S2 ectodomain; and d) the candidate ERGIC retention signal and potentially palmitoylated cysteine residues in the S2 endodomain (Fig. 1). We examined S gp expression, processing, subunit association, glycosylation, and incorporation into lentivirus, vesicular stomatitis virus (VSV), and SARS-CoV-2 virus-like particles (VLPs). We evaluated the infectivity, stability and sensitivity to inhibition of viruses pseudotyped with the S gp variants. We measured the ability of the S gp variants to mediate cell-cell fusion to form lethal syncytia, and found that TMPRSS2 and matrix metalloproteases can contribute to the efficiency of this process. Matrix metalloprotease inhibitors blocked S-mediated cell-cell fusion in TMPRSS2^**—**^cells, and synergized with a TMPRSS2 inhibitor in TMPRSS2^+^ cells. Our results provide insights into the evolutionary features of the SARS-CoV-2 S gp, as well as host factors, that contribute to viral infectivity and pathogenicity.

**FIG 1.**
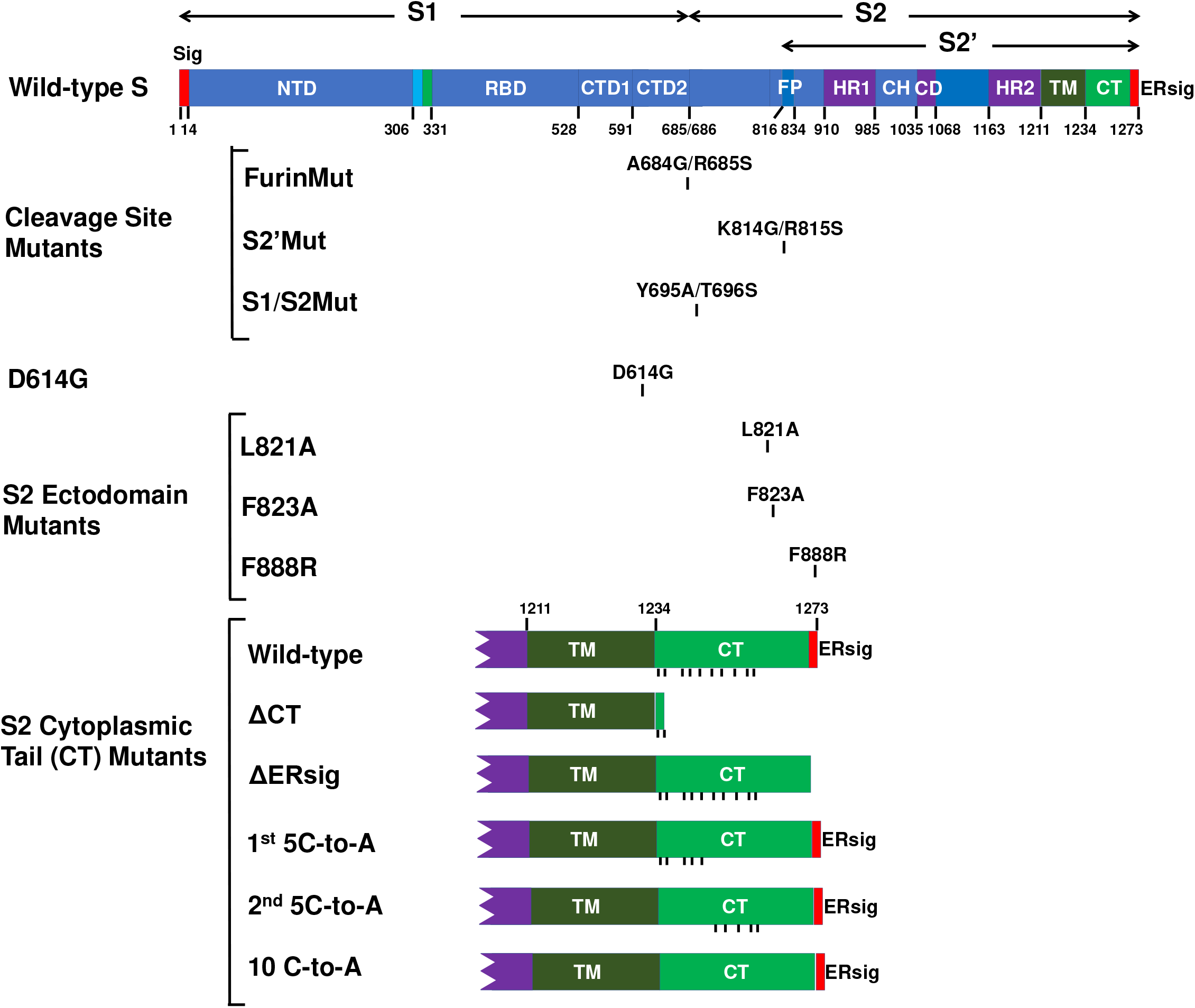
SARS-CoV-2 spike (S) glycoprotein mutants. A schematic representation of the SARS-CoV-2 S gp is shown, with the boundaries of the mature S1, S2 and S2′ glycoproteins indicated. The S gp regions include the signal peptide (Sig), the N-terminal domain (NTD), receptor-binding domain (RBD), C-terminal domains (CTD1 and CTD2), fusion peptide (FP), heptad repeat regions (HR1 and HR2), central helical region (CH), the connector domain (CD), transmembrane region (TM), endodomain/cytoplasmic tail (CT) and endoplasmic reticulum retention signal (ERsig). The changes associated with the S gp mutants studied here are shown. For the S2 cytoplasmic tail mutants, the C-termini of the wild-type and mutant S glycoproteins are depicted, with the positions of cysteine residues indicated by vertical tick marks.

## RESULTS

### Properties of the S gp from the prototypic wild-type SARS-CoV-2 strain

As a reference for comparison with S gp mutants, we evaluated the properties of the wild-type S gp (with Asp 614) derived from the prototypic SARS-CoV-2 strain responsible for the initial outbreak in Wuhan, Hubei province, China. The wild-type S gp was expressed in 293T cells alone or in combination with HIV-1 Gag/protease, which promotes the budding of lentivirus VLPs. The S gp precursor as well as the mature S1 and S2 glycoproteins could be detected in the cell lysates, on the cell surface and on lentivirus VLPs (Fig. 2). The S1 gp was shed into the medium of cells expressing the wild-type S gp alone (Fig. 3A). Very low levels of S gp were detected in the particles prepared from the culture medium of cells expressing the wild-type S gp without HIV-1 Gag, indicating that the vast majority of the S gp pelleted in the presence of HIV-1 Gag is in lentivirus VLPs and not in extracellular vesicles (Fig. 2 and 3B). We confirmed the presence of VLPs of the expected size and morphology decorated with S glycoproteins by electron microscopy with a gold-labeled S-reactive convalescent serum (data not shown). A higher ratio of cleaved:uncleaved S gp was present on lentivirus VLPs than in the cell lysates or on the surface of S gp-expressing cells, as seen previously (32,38) (Fig. 2 and 3B). Selective incorporation of the cleaved S glycoproteins was also observed in VSV pseudotypes (Fig. 4). The uncleaved wild-type S gp in cell lysates and on the cell surface was modified mainly by high-mannose glycans, whereas the cleaved S1 gp contained complex carbohydrates (Fig. 3C). Both the cleaved (major) fraction and uncleaved (minor) fraction of the wild-type S gp incorporated into lentivirus VLPs were modified by complex glycans, indicating passage through the Golgi (51). This result is similar to our finding that HIV-1 envelope glycoproteins can be transported to the cell surface by Golgi and Golgi-bypass pathways, but only those passing through the Golgi are incorporated into HIV-1 virions (Zhang et al., unpublished observations).

**FIG 2.**
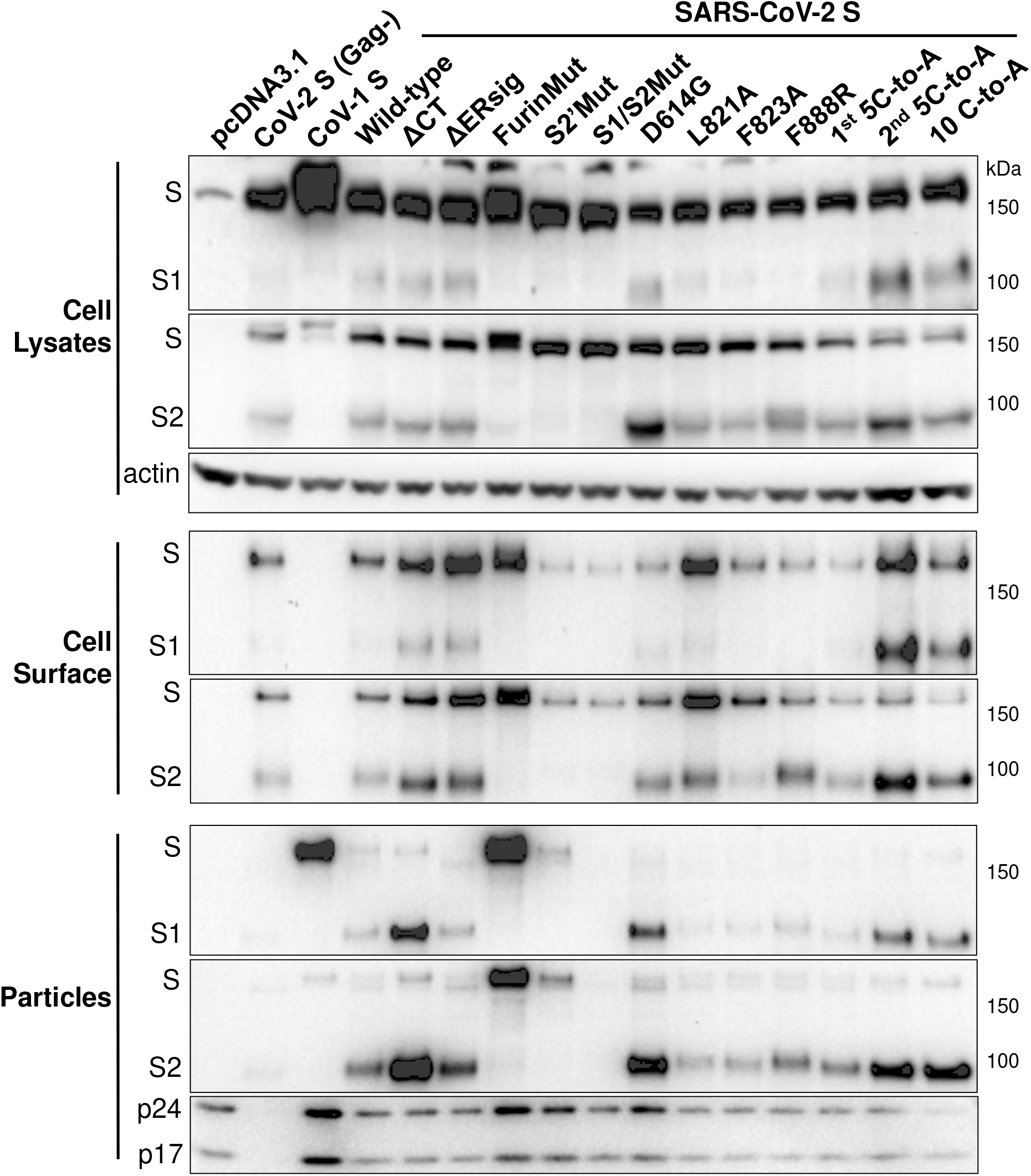
Expression and processing of the SARS-CoV-2 S gp variants. 293T cells were cotransfected with a plasmid encoding HIV-1 Gag/protease and either pcDNA3.1 or plasmids expressing SARS-CoV-1 S gp or wild-type or mutant SARS-CoV-2 S glycoproteins. In Lane 2 (CoV-2 S (Gag-)), 293T cells express the wild-type SARS-CoV-2 S gp without HIV-1 Gag. Cell lysates and VLPs were Western blotted for the S1 and S2 glycoproteins. Cell-surface S glycoproteins were precipitated by convalescent serum NYP01 and then Western blotted for the S1 and S2 glycoproteins (Note that NYP01 does not recognize the SARS-CoV-1 S gp). Cell lysates were Western blotted for actin, and VLPs for HIV-1 p24 and p17 Gag proteins. The results shown are representative of those obtained in at least two independent experiments.

**FIG 3.**
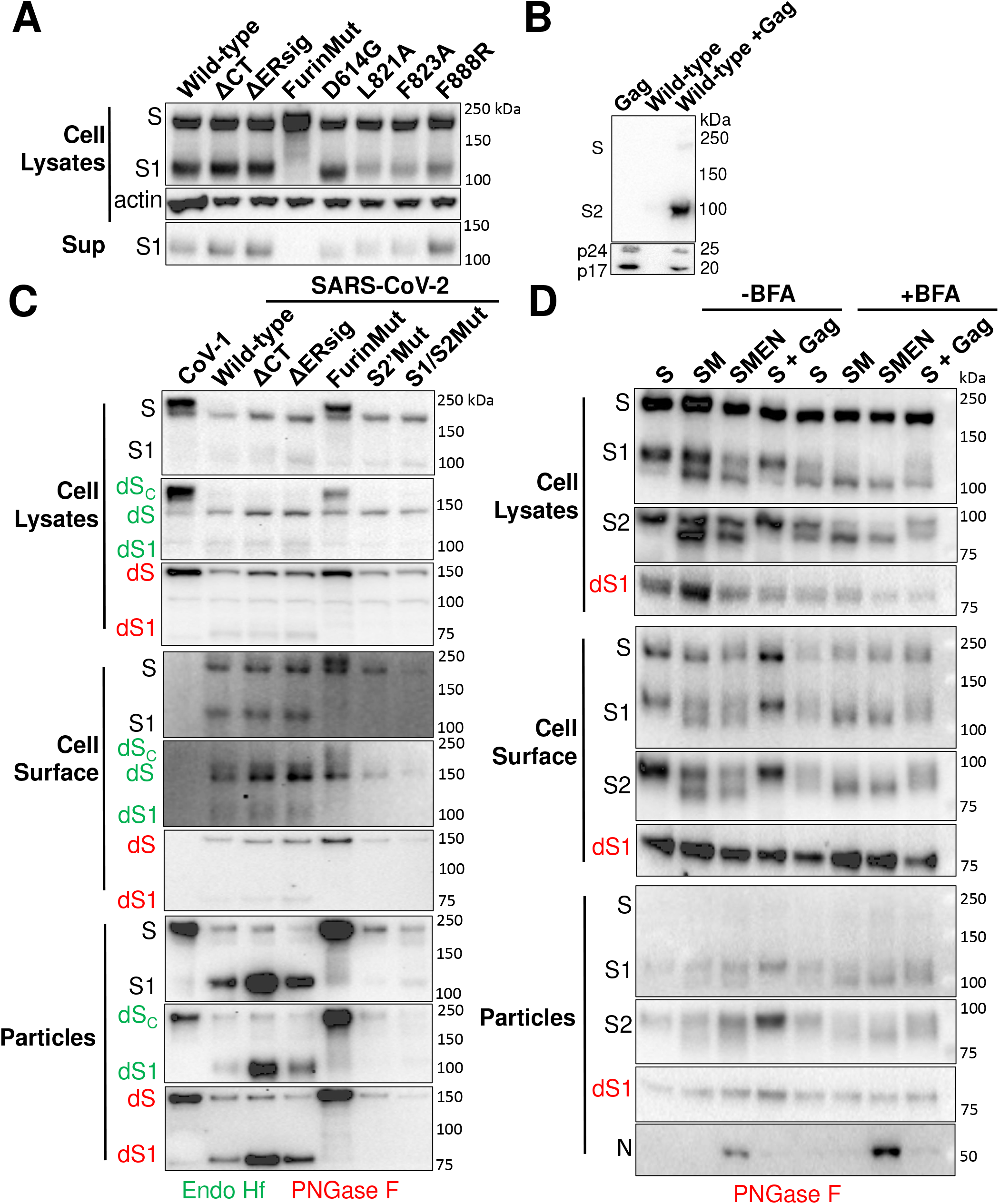
Subunit association, VLP incorporation and glycosylation of S gp variants. (A) 293T cells were transfected with plasmids expressing the indicated SARS-CoV-2 S gp variants. Forty-eight hours later, cell supernatants were filtered (0.45-μm) and precipitated with the convalescent NYP01 serum. The precipitated proteins (Sup, lower panel) and cell lysates (upper panel) were Western blotted with a mouse antibody against S1. Cell lysates were also Western blotted with an antibody against actin. (B) 293T cells were transfected with plasmids expressing HIV-1 Gag/protease and wild-type S gp, separately or together. Two days after transfection, the cell supernatants were cleared by low-speed centrifugation, filtered (0.45-μm) and centrifuged at 14,000 x g for one hour. The pellets were Western blotted for S2 (upper panel) or Gag (lower panel). (C) 293T cells expressing the HIV-1 Gag/protease and the SARS-CoV-1 (Lane 1) or wild-type or mutant SARS-CoV-2 S glycoproteins were used to prepare cell lysates or VLPs. Cell-surface proteins were precipitated by the convalescent serum NYP01. The samples were either mock-treated or treated with Endoglycosidase Hf (green) or PNGase F (red) and then Western blotted for the S1 gp. (D) 293T cells expressing the wild-type SARS-CoV-2 S gp alone or the S gp with the SARS-CoV-2 M or M, E and N proteins or with HIV-1 Gag/protease were treated with Brefeldin A (+BFA) or mock-treated (-BFA) six hours after transfection. The cells were then used to prepare cell lysates, VLPs or cell-surface proteins, as above. The samples were either mock-treated or treated with PNGase F (red) and then Western blotted for the S1 and S2 glycoproteins. The gels in panels A, C and D are representative of those obtained in two independent experiments.

**FIG 4.**
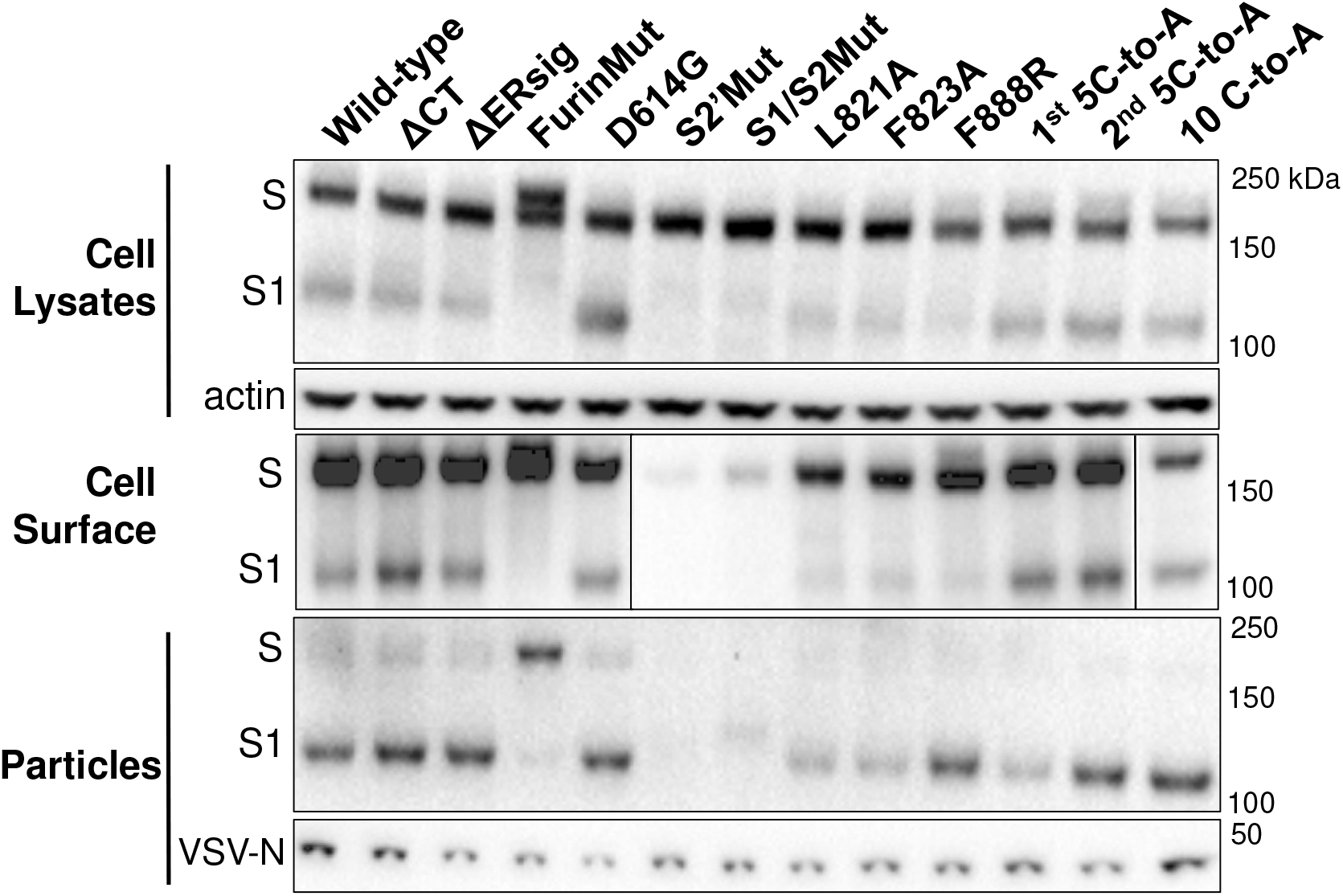
Incorporation of SARS-CoV-2 S gp variants in VSV vectors. 293T cells expressing the indicated SARS-CoV-2 S gp variants and transduced with a recombinant VSVΔG vector were used to analyze the S glycoproteins in cell lysates, on the cell surface and on VLPs. The S glycoproteins were analyzed as described in the Fig. 2 legend except that, in this case, a mouse antibody against S1 was used for Western blotting. The VLP samples were also blotted with an anti-VSV-N antibody. The results shown are representative of those obtained in two independent experiments.

To examine the proteolytic processing and glycosylation of the wild-type S gp in the context of cells producing SARS-CoV-2 VLPs, we coexpressed the M protein or the M, E and N proteins. Expression of these proteins has been shown to result in VLP assembly and budding for the related SARS-CoV-1 (52–56). When the SARS-CoV-2 M protein was coexpressed with the S gp, faster-migrating forms of the S1 and S2 glycoproteins were evident in the cell lysates, on the cell surface and on VLPs (Fig. 3D). In cells treated with Brefeldin A, an inhibitor of anterograde transport through the Golgi (57–59), these faster-migrating forms predominated in M-expressing cells, but were also evident even in cells without M. Sensitivity to PNGase F and Endoglycosidase H identified these S1/S2 species as hypoglycosylated glycoforms modified by high-mannose and/or hybrid carbohydrates (Fig. 3D and data not shown). M coexpression apparently leads to retention of some S gp in early Golgi compartments, where proteolytic cleavage but not complex carbohydrate addition occurs.

To study cytopathic effects mediated by the SARS-CoV-2 S gp, we established the 293T-S and 293T-S-ACE2 cell lines, both of which express the wild-type S gp under the control of a tetracycline-regulated promoter. In addition, the 293T-S-ACE2 cells constitutively express ACE2. Both cell lines propagated efficiently in the absence of doxycycline, a tetracycline analogue (Fig. 5A). 293T-S cells grew nearly as well in the presence of doxycycline as in the absence of the compound. By contrast, doxycycline-induced S gp expression in the 293T-S-ACE2 cells resulted in dramatic cell-cell fusion and cell death. Thus, the coexpression of the SARS-CoV-2 S gp and human ACE2 led to significant cytopathic effects. Similarly, transient expression of the wild-type SARS-CoV-2 S gp in 293T-ACE2 cells resulted in the formation of massive syncytia (Fig. 5B and 6). In the same assay, 293T-ACE2 cells expressing the SARS-CoV-1 S gp did not form syncytia. To quantify the amount of S-mediated cell-cell fusion, we adapted the alpha-complementation assay previously used to measure cell-cell fusion mediated by the HIV-1 envelope glycoproteins (60). Effector cells expressing the wild-type SARS-CoV-2 S gp yielded a signal with ACE2-expressing target cells in the alpha-complementation assay that was approximately 230-fold above that seen for the negative control plasmid (data not shown). These results demonstrate that cells expressing the SARS-CoV-2 S gp fuse efficiently with cells expressing human ACE2.

**FIG 5.**
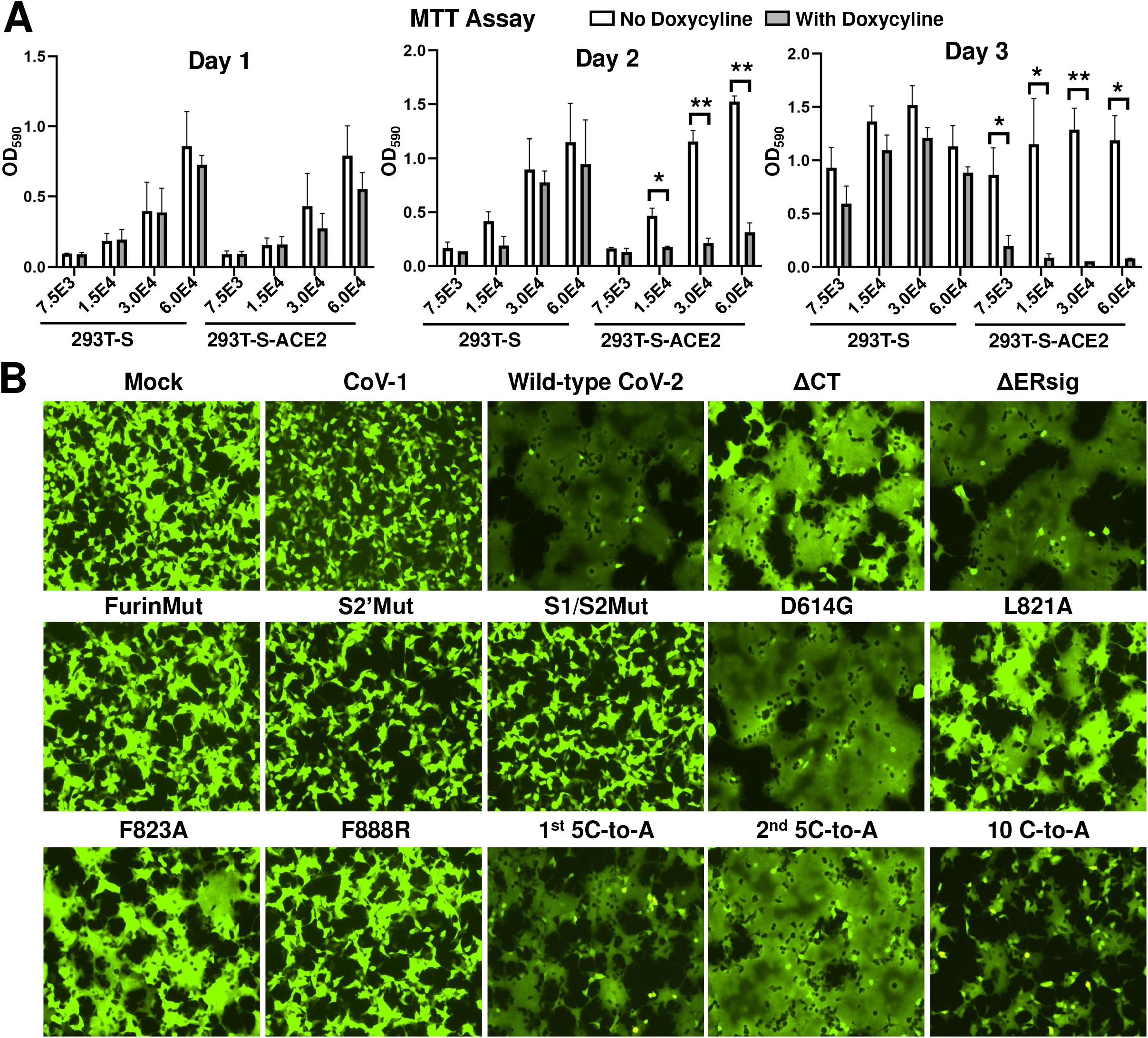
Cytopathic effects mediated by the SARS-CoV-2 S gp variants. (A) The indicated number of 293T-S cells or 293T-S-ACE2 cells, both of which inducibly express the wild-type SARS-CoV-2 S gp under the control of a tetracycline-regulated promoter, were plated in medium with or without doxycycline. On the indicated days after induction with doxycycline, the cells were evaluated by the MTT assay. (B) 293T-ACE2 cells were transfected either with a plasmid expressing enhanced green fluorescent protein (eGFP) alone (Mock) or with the eGFP-expressing plasmid and a plasmid expressing SARS-CoV-1 S gp or wild-type or mutant SARS-CoV-2 S glycoproteins. After 24 hours, the cells were examined under a fluorescent microscope. The results shown are representative of those obtained in two independent experiments.

**FIG 6.**
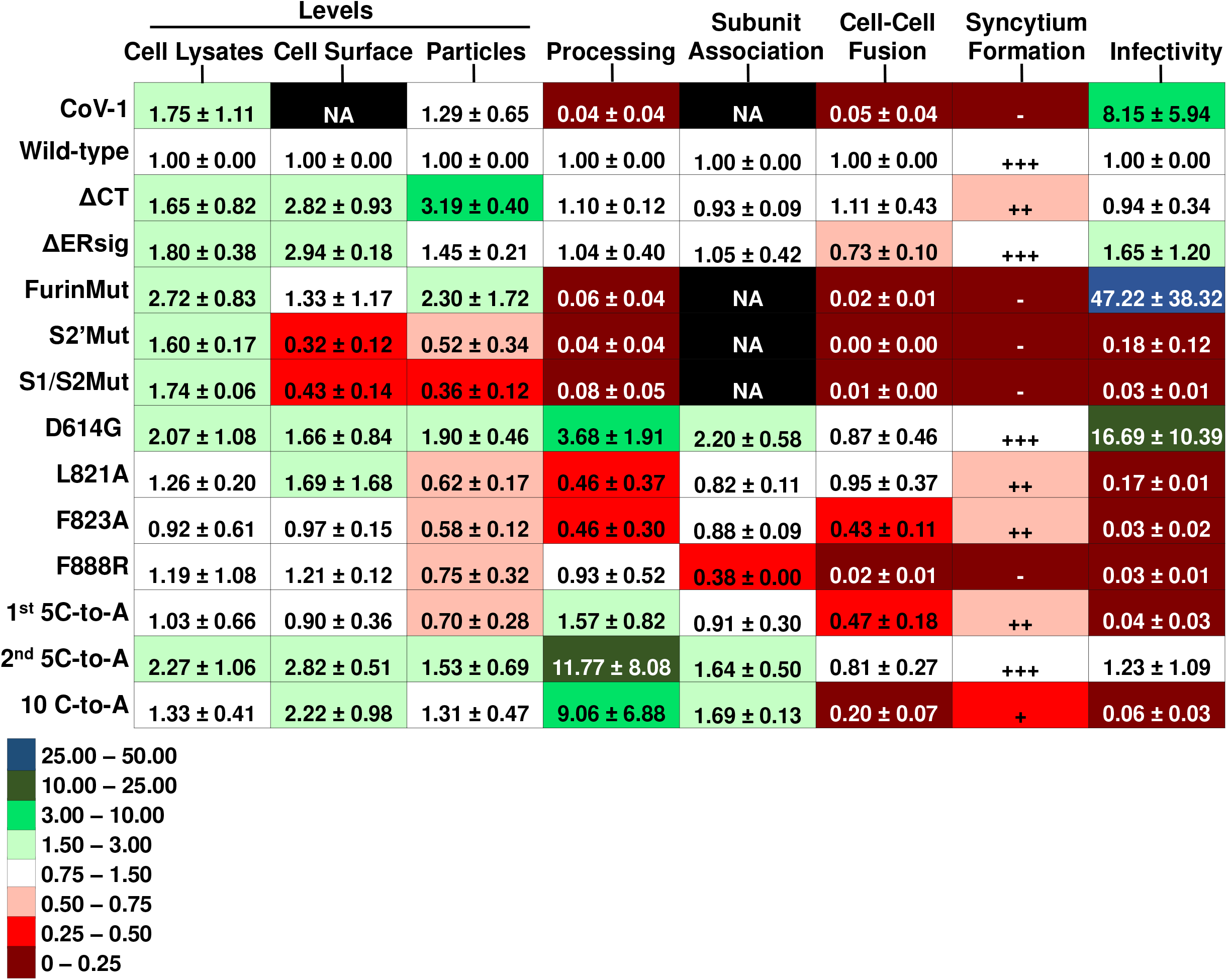
Phenotypes of SARS-CoV-2 S gp variants. Using Western blots similar to that shown in Fig. 2, the total levels of S gp (uncleaved S + S1 + S2) in cell lysates and VLPs and on the surface of 293T cells expressing HIV-1 Gag/protease and the indicated S variant were calculated. The values were normalized to those observed for the wild-type SARS-CoV-2 S gp. The processing index represents the product of the S1/uncleaved S and S2/uncleaved S ratios for each S variant, relative to that of the wild-type SARS-CoV-2 S gp. The association of the S1 and S2 subunits was assessed by measuring the ratio of S1 in lysates to S1 in the supernatants of 293T cells transfected with plasmids expressing the S gp variant alone; the lysate/supernatant S1 ratio for each S gp variant was then normalized to that observed for the wild-type SARS-CoV-2 S gp. Cell-cell fusion represents the activity observed in the alpha-complementation assay for effector cells expressing the indicated S gp variant and ACE2-expressing 293T target cells, relative to the activity observed for the wild-type SARS-CoV-2 S gp. Syncytia were scored by size visually from experiments similar to that shown in Fig. 5B. The infectivity of lentiviruses pseudotyped with the S gp variants, relative to that observed for a virus with the wild-type SARS-CoV-2 S gp, was assessed on 293T-ACE2 target cells. The means and standard deviations derived from at least two independent experiments are shown. NA – not applicable.

To evaluate the ability of the SARS-CoV-2 S gp to mediate virus entry, we measured the single-round infection of target cells by recombinant HIV-1 and VSV vectors pseudotyped with the wild-type S gp. Although the VSV(S) pseudotypes exhibited higher infectivity than HIV-1(S) pseudotypes, infection of 293T-ACE2 cells by both S-pseudotyped vectors was at least 60-fold above the background seen for viruses without envelope glycoproteins (data not shown). Consistent with previous findings (38,61), expression of TMPRSS2 in the 293T-ACE2 target cells increased the efficiency of virus infection mediated by the wild-type SARS-CoV-2 S gp (Fig. 7A).

**FIG 7.**
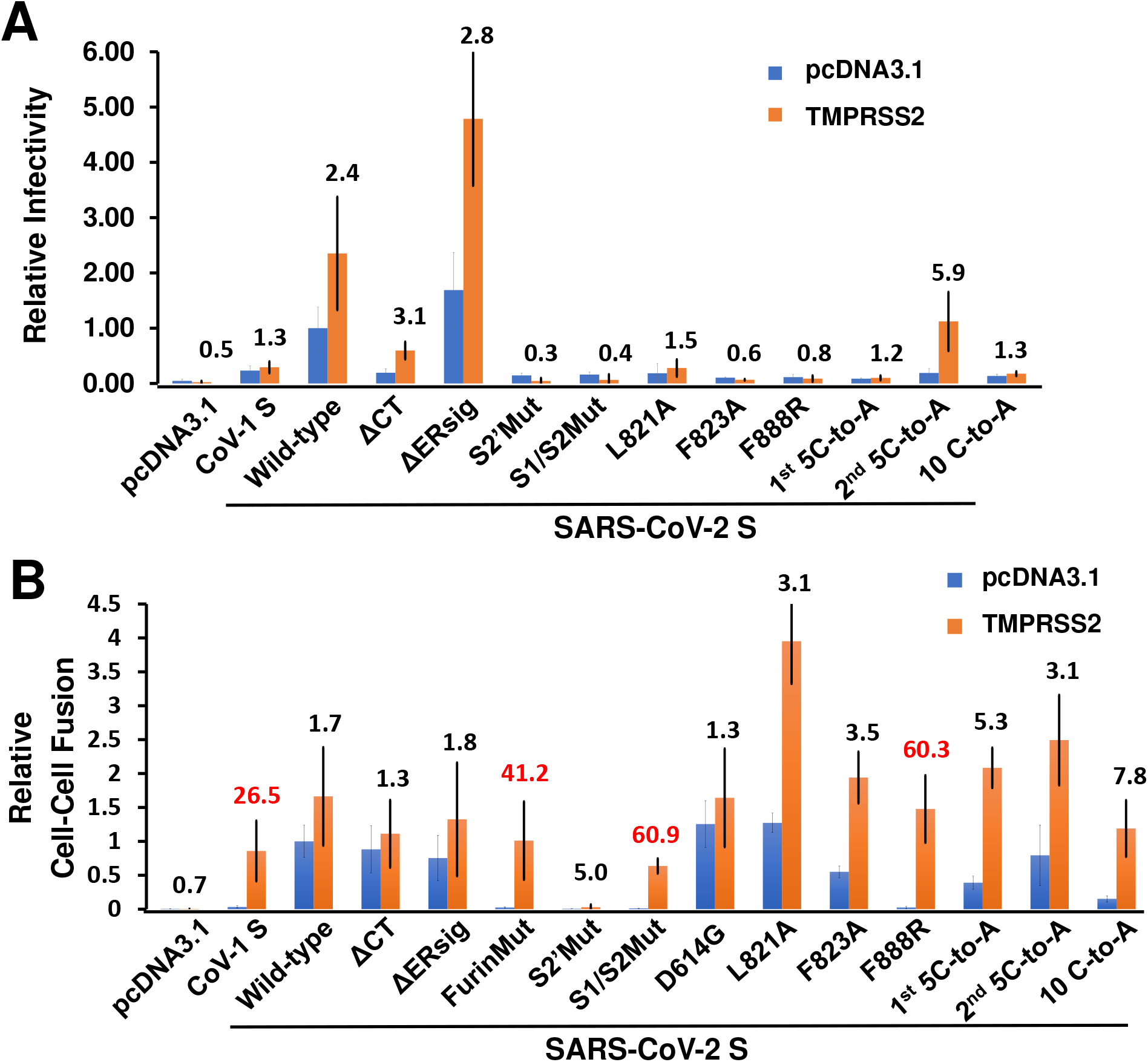
Effect of TMPRSS2 expression in target cells on SARS-CoV-2 S gp-mediated infectivity and cell-cell fusion. (A) The infectivity of VSV vectors pseudotyped with the indicated S gp variants for 293T-ACE2 target cells transfected either with pcDNA3.1 or a TMPRSS2-expressing plasmid is shown, relative to the value for the wild-type S gp in 293T-ACE2 target cells. The fold increase in infection associated with TMPRSS2 expression in the target cells is indicated above the bar graphs. (B) Effector cells expressing alpha-gal and different SARS-CoV-2 S gp variants were cocultivated for four hours at 37°C with 293T cells expressing omega-gal and either ACE2 + pcDNA3.1 or ACE2 + TMPRSS2. The beta-galactosidase activity is shown for each S gp variant, relative to that seen with effector cells expressing the wild-type S gp and target cells expressing ACE2. The fold increase associated with TMPRSS2 expression in the target cells is indicated above the bar graphs; the greatest increases are highlighted in red. The results shown are representative of those obtained in two independent experiments. The means and standard deviations from triplicate wells are shown.

### The furin cleavage site reduces virus infectivity, stability and sensitivity to ACE2 and neutralizing antibodies, but enhances S gp syncytium-forming ability

At some point during its evolution from an ancestral bat coronavirus, the SARS-CoV-2 S gp acquired a multibasic cleavage site for furin-like proteases at the S1/S2 junction (9,15,41,42). To evaluate the impact of this evolutionary change, we altered the two residues immediately N-terminal to the proposed furin cleavage site (FurinMut in Fig. 1). Compared with the wild-type SARS-CoV-2 S gp, the FurinMut S gp was expressed at higher levels in 293T cells and was efficiently incorporated into lentivirus VLPs, but was inefficiently processed into S1 and S2 glycoproteins (Fig. 2 and 3A). Two glycoforms of the uncleaved FurinMut, one with only high-mannose glycans and the other with complex glycans, were present in cell lysates and on the cell surface; however, only the glycoform modified by complex carbohydrates and presumably transported through the Golgi was found in lentivirus VLPs (Fig. 3C).

In contrast to the wild-type S gp, FurinMut did not mediate syncytium formation or induce cell-cell fusion in the alpha-complementation assay (Fig. 5B and 6). However, the cell-fusing activity of the FurinMut S gp was dramatically increased by coexpression of TMPRSS2 and ACE2 in 293T target cells (Fig. 7B). The infectivity of the FurinMut-pseudotyped virus was 7-50 times higher than that of viruses pseudotyped with the wild-type S gp; the infectivity difference between viruses pseudotyped with FurinMut and wild-type S glycoproteins was greater for HIV-1 pseudotypes than for VSV pseudotypes (Fig. 6 and 8A). The infectivity of the FurinMut virus incubated on ice was more stable than that of the wild-type S virus (Fig. 8B). FurinMut viruses were more sensitive to inhibition by soluble ACE2 and convalescent sera than wild-type viruses (Fig. 8C). Thus, the acquisition of the furin cleavage site by the SARS-CoV-2 S gp decreased virus stability, infectivity and sensitivity to soluble ACE2 and neutralizing antibodies, but greatly enhanced the ability to form syncytia.

**FIG 8.**
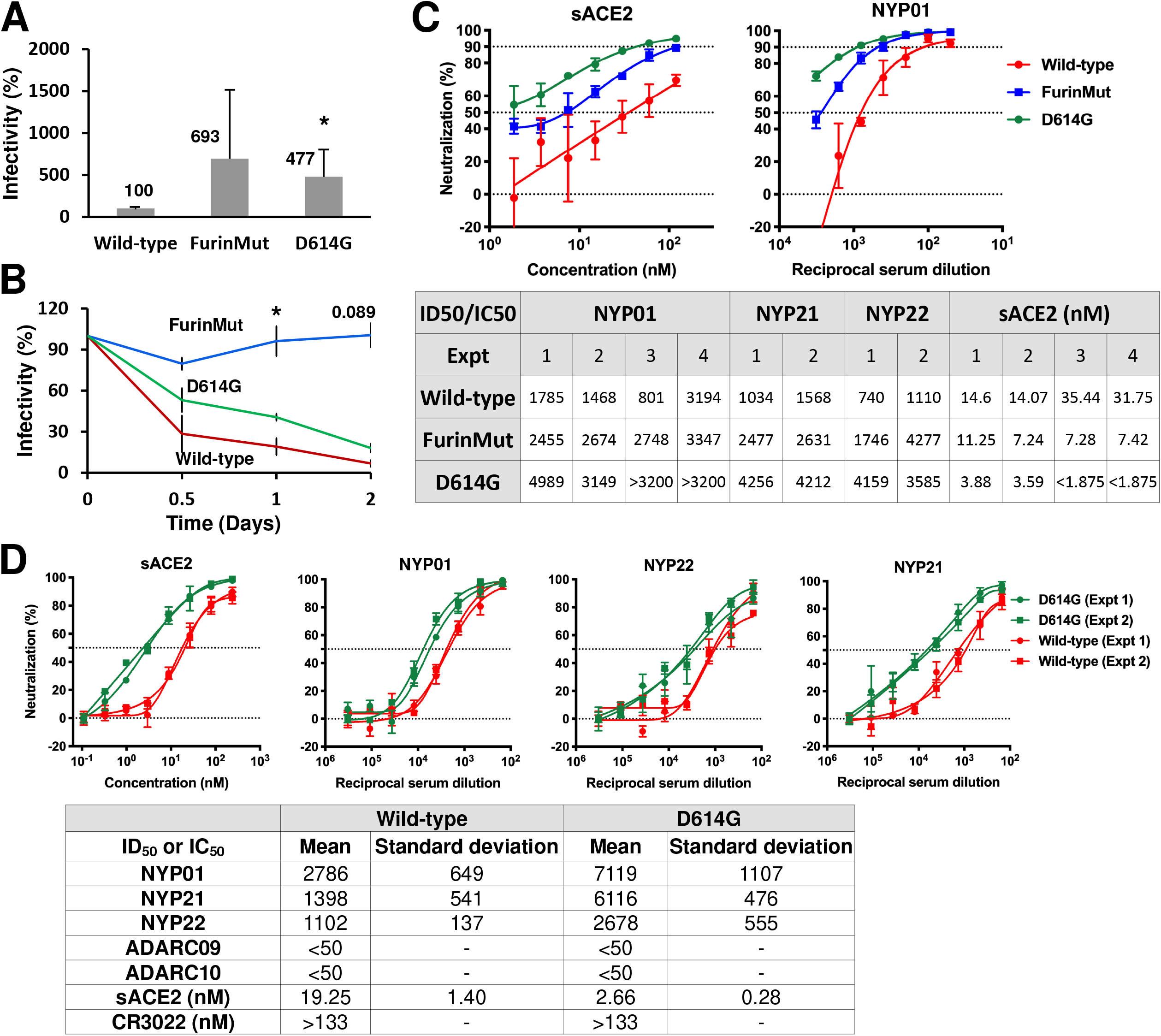
The D614G and FurinMut changes influence virus infectivity, cold sensitivity and sensitivity to soluble ACE2 and neutralizing antisera. (A) The infectivity of VSV vectors pseudotyped with the indicated SARS-CoV-2 S gp variants for 293T-ACE2 target cells is shown, relative to the value seen for the wild-type S gp. The means and standard deviations derived from at least 4 independent experiments are shown. (B) VSV vectors pseudotyped with the indicated SARS-CoV-2 S glycoproteins were incubated on ice for the indicated times, after which the infectivity on 293T-ACE2 cells was assessed. The measured infectivities were normalized to that observed for each S gp variant at Time 0. (C-D) VSV vectors pseudotyped with the wild-type, FurinMut or D614G S gp variants were used to infect 293T-ACE2 cells following incubation with different dilutions/concentrations of NYP01, NYP21 and NYP22 convalescent sera or soluble (sACE2). The dilutions/concentrations of sera and ACE2 required to inhibit 50% of infection are shown. The results of independent experiments are shown. In panel D, the means and standard deviations derived from 3-4 independent experiments are reported. ADARC09 and ADARC10 are control sera from individuals not infected by SARS-CoV-2. Statistical significance was evaluated using Student’s t-test. *, P < 0.05.

### Alteration of other potential cleavage sites reduces S gp processing and function

The SARS-CoV-1 S gp lacks a favorable cleavage site for furin-like proteases, but during infection of a target cell is thought to be cleaved at a nearby secondary site and at the S2′ site by cellular proteases (Cathepsin L, TMPRSS2) (62–64). Alteration of the secondary cleavage site near the S1/S2 junction (S1/S2Mut) in the SARS-CoV-2 S gp resulted in complete lack of proteolytic processing, inefficient incorporation into lentivirus VLPs, and loss of function in cell-cell fusion and infectivity assays (Fig. 2, 3C, 5B and 6). Alteration of the S2′ site (S2′Mut) also led to lack of S processing, although low levels of VLP incorporation and infectivity were detected. The S2′Mut gp in the VLPs was Endoglycosidase H-resistant, suggesting that it is modified by complex carbohydrates (Fig. 3C). The S2′Mut gp did not detectably mediate cell-cell fusion (Fig. 6). Thus, despite the presence of the furin cleavage site in these mutants, proteolytic processing did not occur. Both mutants exhibited severe decreases in infectivity and cell-cell fusion. However, coexpression of TMPRSS2 in the ACE2-expressing target cells enhanced cell-cell fusion by the S1/S2Mut gp but not by S2′Mut gp (Fig. 7B).

### The D614G change in the predominant SARS-CoV-2 strain increases S1-trimer association, virus infectivity and sensitivity to soluble ACE2 and neutralizing antisera

The change in Asp 614 to a glycine residue (D614G) is found in the predominant emerging SARS-CoV-2 strains worldwide (43,44). The D614G S gp was cleaved slightly more efficiently than the wild-type S gp, but shed less S1 into the medium of expressing cells (Fig. 2, 3A and 6). The efficiencies of cell-cell fusion mediated by the D614G and wild-type S glycoproteins were comparable (Fig. 5B and 6). Both lentivirus and VSV vectors pseudotyped with the D614G S gp infected cells 4-16-fold more efficiently than viruses with the wild-type S gp (Fig. 6 and 8A). The stabilities of the viruses pseudotyped with D614G and wild-type S glycoproteins on ice were comparable (Fig. 8B). Importantly, the viruses with D614G S gp were approximately 7-fold more sensitive to soluble ACE2 and 2-5-fold more sensitive to neutralizing antisera than viruses with wild-type S gp (Fig. 8D). Soluble ACE2 bound and induced the shedding of S1 from D614G VLPs more efficiently than from VLPs with the wild-type S gp (Fig. 9A and B). The lentivirus VLPs with D614G S gp exhibited a significantly greater association of the S1 subunit with the trimer (half-life greater than 5 days at 37°C) compared with viruses with wild-type S gp (half-life 2-3 days at 37°C) (Fig. 9C). S1 association with detergent-solubilized S gp trimers was greater for D614G than for wild-type S over a range of temperatures from 4-37°C (Fig. 9D). Thus, the D614G change enhances virus infectivity, responsiveness to ACE2 and S1 association with the trimeric spike.

**FIG 9.**
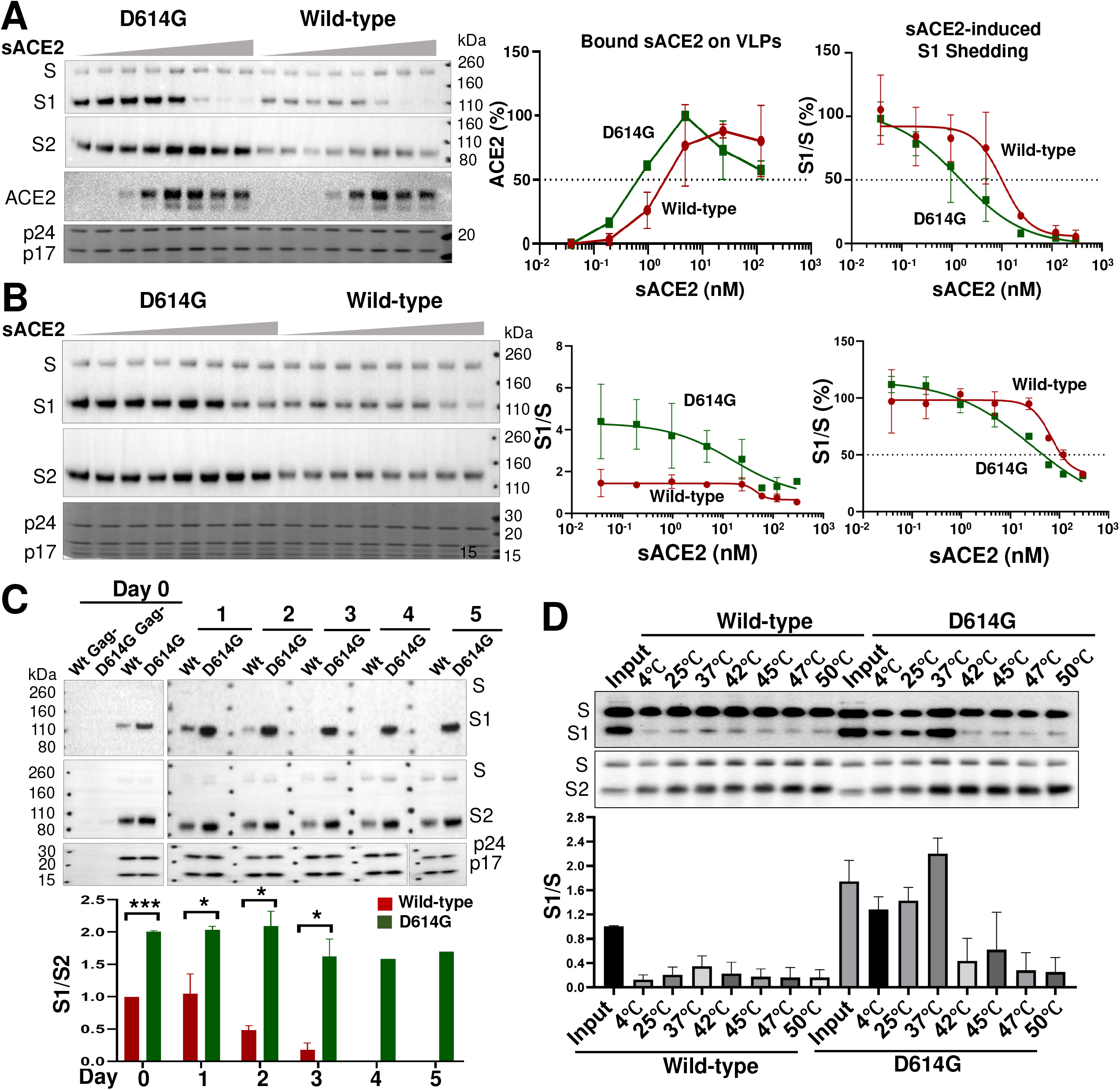
D614G mutation increases S gp sensitivity to soluble ACE2 and S1-trimer association. (A) HIV-1 VLPs pseudotyped with the wild-type or D614G S glycoproteins were incubated with various concentrations (0 - 300 nM) of soluble ACE2 (sACE2) for 1 hr on ice. The VLPs were pelleted and lysed. VLP lysates were Western blotted for S1, S2 and ACE2. HIV-1 p24 and p17 Gag proteins in the VLP lysates were detected by Coomasie blue staining. (B) HIV-1 VLPs pseudotyped with the wild-type or D614G glycoproteins were incubated with the indicated concentrations of sACE2 for 1 hr at 37°C. The VLPs were pelleted and lysed. VLP lysates were Western blotted for S1 and S2, and HIV-1 p24 and p17 Gag proteins were detected by staining with Coomassie blue. The S1/S ratio as a function of sACE2 concentration is shown in the graphs in the middle and right panels. In the graph on the right, the S1/S ratios are normalized to that seen for the wild-type S gp in the absence of sACE2, which is set at 100%. (C) HIV-1 VLPs pseudotyped with the wild-type or D614G S glycoproteins were incubated at 37°C for the indicated number of days, after which the VLPs were pelleted, lysed and Western blotted. As negative controls, supernatants from 293T cells expressing the S glycoproteins without HIV-1 Gag were processed in parallel (lanes 1 and 2, Gag-). (D) Lysates from 293T cells transiently expressing the His_6_-tagged wild-type or D614G glycoproteins were incubated at the indicated temperatures for 1 hr. The S glycoproteins were then precipitated by Ni-NTA resin and Western blotted. The blots shown are representative of those obtained in two independent experiments and the graphs show the means and standard deviations from two independent experiments. Statistical significance was evaluated using Student’s t-test. *, P < 0.05; **, P < 0.01; ***, P < 0.001.

### Changes in the S2 fusion peptide decrease infectivity

The putative fusion peptide of the SARS-CoV-2 S2 glycoprotein (residues 816-834) has been identified by analogy with the SARS-CoV-1 S gp. Changes in the fusion peptide and a more C-terminal S2 region in SARS-CoV-1 S gp have been suggested to result in fusion-defective mutants (65–67). We introduced analogous changes into the putative fusion peptide of the SARS-CoV-2 S gp (L821A and F823A), and also made a change (F888R) in the downstream region implicated in SARS-CoV-1 S gp function. The L821A and F823A mutants were processed slightly less efficiently than the wild-type S gp. Compared with the wild-type S gp, the F888R mutant exhibited a lower ratio of the S1 gp in cell lysates relative to the S1 gp in cell supernatants, suggesting a decrease in the association of S1 with the S trimer (Fig. 3A and 6). Modest decreases in the level of S glycoproteins on lentivirus VLPs were observed for all three mutants. L821A and F823A retained the ability to mediate cell-cell fusion, although the syncytia formed were smaller than those induced by the wild-type S gp (Fig. 5B and 6). F888R was severely compromised in the ability to mediate cell-cell fusion; however, this ability was recovered when TMPRSS2 was coexpressed with ACE2 in the target cells (Fig. 7B). The infectivity of viruses pseudotyped with these three mutants was greatly decreased, relative to viruses with wild-type S gp (Fig. 6 and 7A). In summary, these S2 ectodomain changes exert pleiotropic effects, ultimately compromising virus infectivity.

### Palmitoylated membrane-proximal cysteines in the S2 endodomain contribute to virus infectivity

The S2 endodomain of SARS-CoV-1 is palmitoylated, contains an ERGIC-retention signal, and contributes to the interaction of S with the M protein during virus assembly (45–50). We altered the highly similar endodomain of the SARS-CoV-2 S gp, which contains ten cysteine residues potentially available for palmitoylation. The endodomain/cytoplasmic tail was deleted in the ΔCT mutant, leaving only the two membrane-proximal cysteine residues (Fig. 1). In the ΔERsig mutant, the putative ERGIC-retention signal at the C terminus of the endodomain was deleted. In three additional mutants, the N-terminal five cysteine residues (1^st^ 5C-to-A), the C-terminal five cysteine residues (2^nd^ 5C-to-A), or all ten cysteine residues (10 C-to-A) were altered to alanine residues.

The ΔCT and ΔERsig mutants were expressed on the cell surface at higher levels than that of the wild-type S gp, and the level of ΔCT on lentivirus VLPs was significantly increased (Fig. 2, 3C and 6). Both ΔCT and ΔERsig mediated syncytium formation and virus infection at levels comparable to those of the wild-type S gp (Fig. 5B and 6). Therefore, except for the two membrane-proximal cysteine residues, the SARS-CoV-2 S2 endodomain is not absolutely required for cell-cell fusion or virus entry.

Proteolytic processing of the 2^nd^ 5C-to-A and 10 C-to-A mutants was more efficient than that of the wild-type S gp (Fig. 2 and 6). Although all three mutants with altered endodomain cysteine residues were expressed on the cell surface and incorporated into VLPs, only the 2^nd^ 5C-to-A mutant supported cell-cell fusion and virus infection at wild-type S levels. Of note, the infectivity of lentiviruses pseudotyped with the 1^st^ 5C-to-A and 10 C-to-A mutants was near the background of the assay. These results implicate the membrane-proximal cysteine residues of the S2 endodomain in the virus entry process.

To evaluate the palmitoylation of the wild-type and mutant S glycoproteins, we used hydroxylamine cleavage and mPEG-maleimide alkylation to mass-tag label the palmitoylated cysteine residues (68). The majority of the wild-type S2 gp was palmitoylated, with different species containing one to four acylated cysteines (Fig. 10). The ΔCT, 1^st^ 5C-to-A and 2^nd^ 5C-to-A mutants consisted of two species, one without palmitoylation and the other with a single palmitoylated cysteine. The ratio of palmitoylated:unmodified S2 glycoprotein was significantly greater for the 2^nd^ 5C-to-A mutant than for the ΔCT and 1^st^ 5C-to-A mutants. The 10 C-to-A mutant was not detectably palmitoylated. Thus, palmitoylation can occur on multiple different cysteine residues in the SARS-CoV-2 S2 endodomain; however, palmitoylation appears to occur more efficiently on the N-terminal endodomain cysteines, which contribute to virus infectivity.

**FIG 10.**
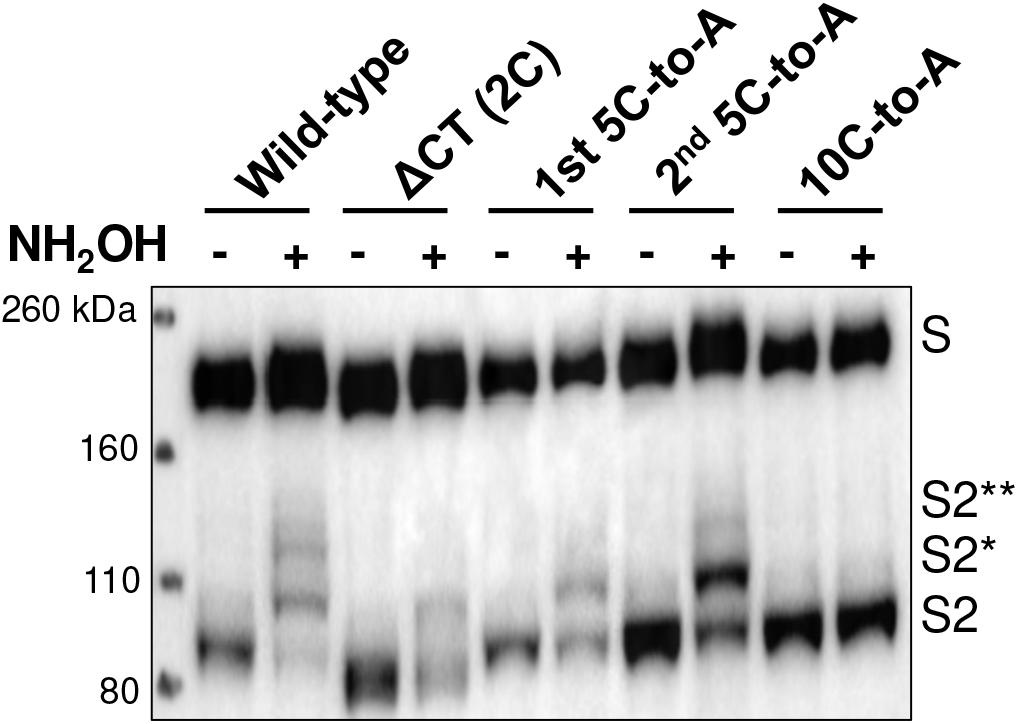
Palmitoylation of the SARS-CoV-2 S2 endodomain/cytoplasmic tail. Lysates prepared from 293T cells expressing the wild-type SARS-CoV-2 S gp or the indicated mutants were subjected to acyl-PEG exchange (68). When NH_2_OH is left out, palmitoylated cysteine residues are not de-acylated and therefore not available for reaction with mPEG-maleimide. The cell lysates were Western blotted with an anti-S2 antibody. The mono-PEGylated (S2*) and di-PEGylated (S2**) species are indicated. Note that the ΔCT mutant retains two membrane-proximal cysteine residues. The results shown are representative of those obtained in two independent experiments.

We also tested whether the effect of M protein coexpression on S glycosylation depended on an intact S endodomain. Surprisingly, M coexpression resulted in the appearance of hypoglycosylated forms of the ΔCT, ΔERsig and 10 C-to-A mutants (data not shown). Thus, an intact or palmitoylated endodomain is not required for M to exert an effect on S glycosylation.

### Metalloprotease inhibitors block S-mediated cell-cell fusion

To gain insight into the host cell factors that might influence our experimental outcomes, we tested several inhibitors in the cell-cell fusion, syncytium formation, and lentivirus pseudotype infection assays. Inhibitors of several proteases potentially involved in S gp activation were tested. E64d, which blocks the activity of cysteine proteases like Cathepsin B/L, partially inhibited the infection of lentiviruses pseudotyped with SARS-CoV-2 S gp at concentrations up to 100 μM (data not shown). E64d had no effect on SARS-CoV-2 S-mediated syncytium formation. In contrast, two broad-spectrum matrix metalloprotease inhibitors, marimastat and ilomastat, inhibited S-mediated cell-cell fusion and syncytium formation (Fig. 11A-C). These inhibitory effects were specific, because neither compound decreased cell-cell fusion or syncytium formation mediated by HIV-1 envelope glycoprotein trimers. Another metalloprotease inhibitor, TAPI-2, also inhibited S-mediated cell-cell fusion, but was less potent and specific than marimastat or ilomastat (data not shown). Cell-cell fusion assays were conducted in which the compounds were incubated either with the S-expressing or ACE2-expressing cells, followed by cocultivation of these cells. The results indicate that the primary inhibitory effect of the metalloprotease inhibitors is exerted on the ACE2-expressing target cells (data not shown). Consistent with this interpretation, no effect of marimastat or ilomastat on the processing of the SARS-CoV-2 S gp was observed (Fig. 11D). None of the metalloprotease inhibitors affected the efficiency of lentivirus infection mediated by the SARS-CoV-2 S gp, at doses up to 100 μM (data not shown). These results suggest that one or more matrix metalloproteases on the surface of the ACE2-expressing target cells contribute to the efficiency of cell-cell fusion mediated by the SARS-CoV-2 S gp. Of interest, marimastat and ilomastat were much less effective at inhibiting S-mediated cell-cell fusion when TMPRSS2 was coexpressed with ACE2 in the target cells (Fig. 12A). Camostat mesylate, an inhibitor of serine proteases like TMPRSS2, when used alone had little or no effect on S-mediated cell-cell fusion, syncytium formation or virus infection of 293T-ACE2 target cells (Fig. 12B, C and data not shown). However, camostat mesylate and marimastat syngergistically inhibited S-mediated syncytium formation, consistent with TMPRSS2 and metalloproteases carrying out redundant functions during the cell-cell fusion process (Fig. 12B and C).

**FIG 11.**
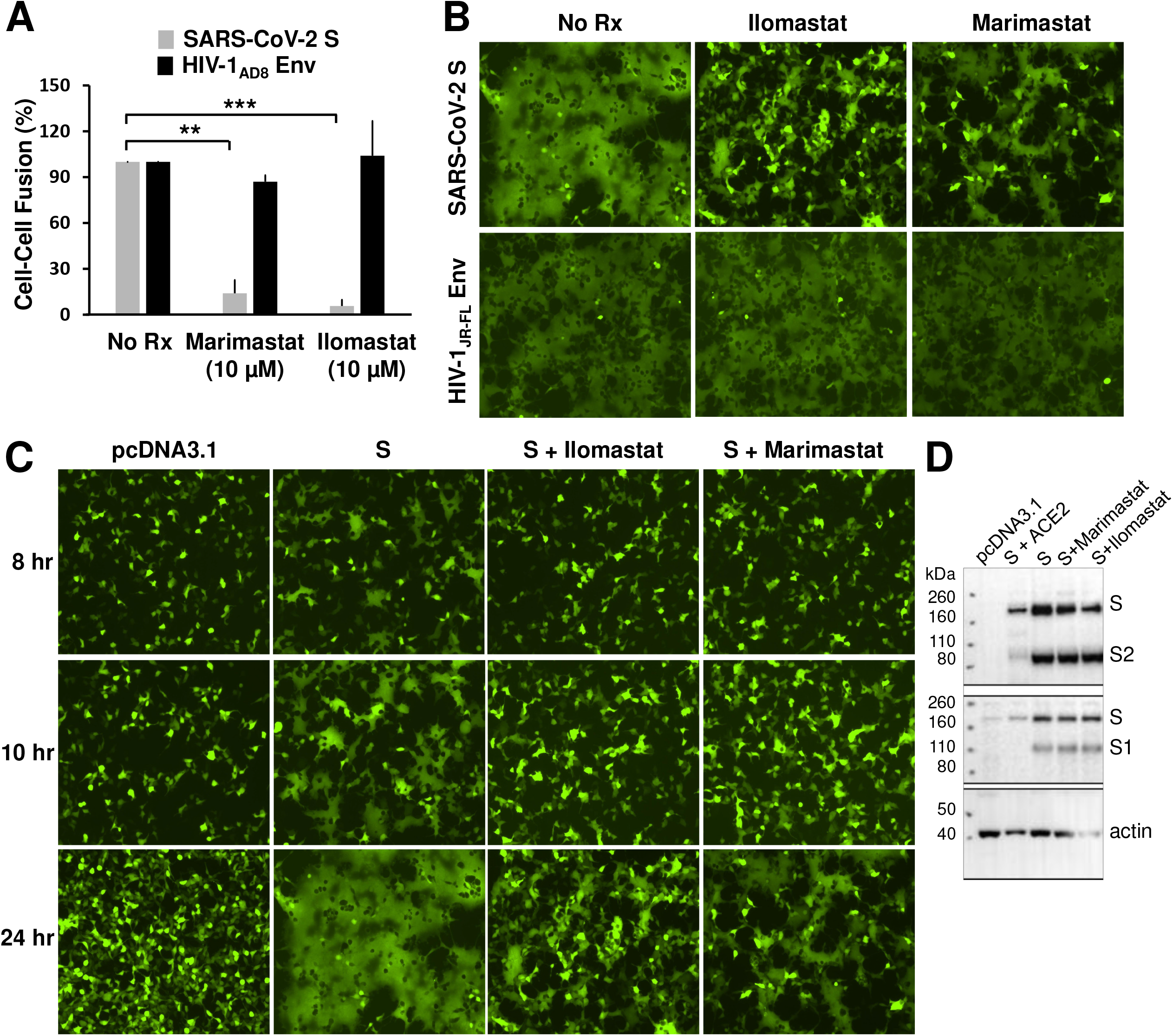
Metalloprotease inhibitors block S-mediated cell-cell fusion and syncytium formation. (A) The effect of marimastat and ilomastat, two matrix metalloprotease inhibitors, on two parallel alpha-complementation assays was tested. In one assay, the effector cells express the wild-type SARS-CoV-2 S gp and 293T cells expressing ACE2 were used as target cells. In the second assay, the effector cells express the HIV-1_AD8_ envelope glycoproteins and the target cells express the CD4 and CCR5 receptors. Target cells were treated with 10 μM inhibitor for 2 hr and then cocultivated with effector cells for 4 hr. The β-galactosidase values were normalized to those seen in the absence of inhibitors. The means and standard deviations from at least two independent experiments are shown. (B) 293T-ACE2 cells were cotransfected with plasmids expressing eGFP and the wild-type SARS-CoV-2 S gp, after which ilomastat (10 μM) or marimastat (10 μM) was added. For the HIV-1 control, 293T cells were transfected with plasmids expressing eGFP, CD4, CCR5 and the HIV-1_JR-FL_ envelope glycoproteins. The cells were imaged 24 hours after transfection. (C) The time course is shown for 293T-ACE2 cells transfected with an eGFP-expressing plasmid and either pcDNA3.1 or a plasmid expressing the wild-type SARS-CoV-2 S gp, in the absence (S) or presence of ilomastat (10 μM) or marimastat (10 μM). (D) 293T cells were transfected with pcDNA3.1, a plasmid expressing the wild-type SARS-CoV-2 S gp alone, or plasmids expressing the wild-type SARS-CoV-2 S gp together with human ACE2. The cells were untreated or treated with marimastat (10 μM) or ilomastat (10 μM). Cell lysates were Western blotted with antibodies against S1 and S2. The results shown in panels B and C are representative of those obtained in two independent experiments. Statistical significance was evaluated using Student’s t-test. *, P < 0.05; **, P < 0.01; ***, P < 0.001.

**FIG 12.**
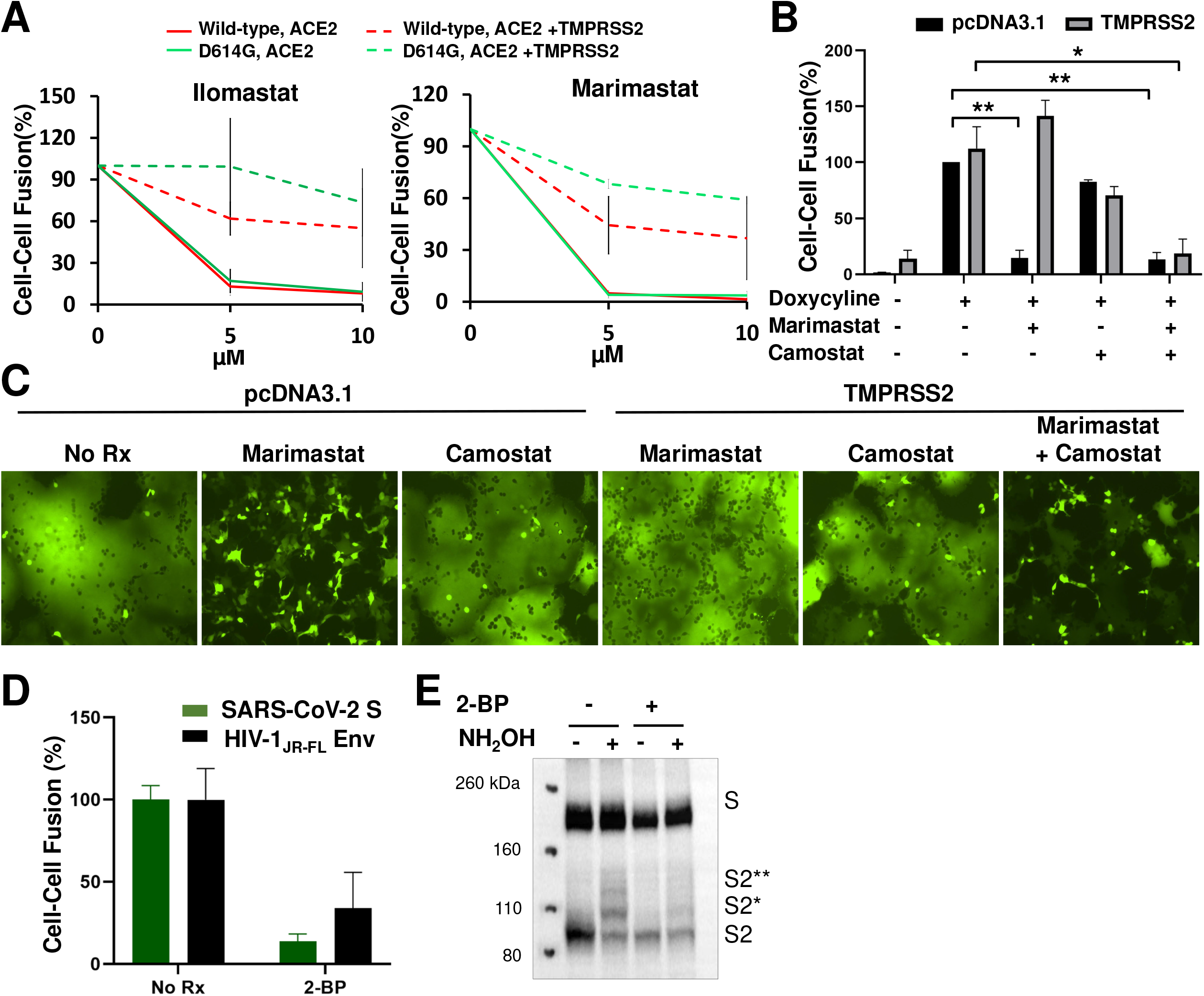
Effect of inhibitors in the absence and presence of TMPRSS2 expression in target cells on SARS-CoV-2 S gp-mediated cell-cell fusion and syncytium formation. (A) In this alpha-complementation assay, 293T target cells expressing ACE2 only or ACE2 + TMPRSS2 were incubated with ilomastat and marimastat at the indicated concentration for 2 hours before cocultivation with effector cells expressing wild-type or D614G S gp for 4 hours. The results with target cells expressing ACE2 are shown as solid lines, and the results with target cells expressing ACE2 and TMPRSS2 are shown as dashed lines. In (B,C), we tested the effects of camostat mesylate (100 μM) and marimastat (10 μM) individually or in combination on cell-cell fusion (B) and syncytium formation (C) in cells expressing ACE2 with or without TMPRSS2 (See Methods). In (B), 293T-S cells inducible with doxycycline were used as effector cells. (D) In the alpha-complementation assays measuring cell-cell fusion mediated by the wild-type SARS-CoV-2 S gp and the HIV-1 envelope glycoproteins, treatment of the effector cells with 100 μM 2-bromopalmitate (2-BP) reduced cell-cell fusion. (E) 293T cells expressing the wild-type SARS-CoV-2 S gp were mock-treated or treated with 100 μM 2-BP. Lysates from these cells were subjected to acyl-PEG exchange as in Fig. 10 and then Western blotted for the S2 gp. Treatment with 2-BP reduces S2 palmitoylation, but also results in decreased S gp processing. The results shown in panels A, C and E are representative of those obtained in two independent experiments. In panels B and D, the means and standard deviations from two independent experiments are shown. Statistical significance was evaluated using Student’s t-test. *, P < 0.05; **, P < 0.01; ***, P < 0.001.

Given the phenotype of the 10 C-to-A mutant, we tested the effects of a palmitoylation inhibitor, 2-bromopalmitate (2-BP) on the function of the wild-type S gp. 2-BP inhibited SARS-CoV-2 S-mediated cell-cell fusion, syncytium formation and virus infection (Fig. 12D and data not shown). We verified that 2-BP treatment reduced palmitoylation of the S2 glycoprotein, but found that 2-BP also reduced the proteolytic processing of the S precursor (Fig. 12E). Cell-cell fusion and syncytium formation mediated by the HIV-1_JR-FL_ envelope glycoprotein were also inhibited by 2-BP (Fig. 12D and data not shown). Treatment of the effector and target cells individually with 2-BP suggested that the main effect of 2-BP is exerted on the cells expressing the SARS-CoV-2 S and HIV-1 envelope glycoproteins. These results indicate that inhibition of palmitoylation can result in blockade of SARS-CoV-2 S gp function, perhaps through indirect effects. For example, furin palmitoylation is required for association of furin with plasma membrane microdomains and processing of some substrates (69).

## DISCUSSION

Many properties contribute to the ability of a virus like SARS-CoV-2 to achieve zoonotic transmission into humans, to spread globally in the human population, to evade host immune systems, and to cause disease. The exposed nature of the spike trimer on the surface of infected cells and viruses renders the S gp particularly subject to evolutionary pressure. Here, we examined several features of the S gp to understand their potential contribution to SARS-CoV-2 infectivity, stability, resistance to neutralization and cytopathic effects.

Conformational changes in Class I viral envelope glycoproteins contribute to virus entry and evasion of host antibody responses (Fig. 13). The pretriggered, “closed” conformation of the SARS-CoV-2 S gp trimer is converted by binding the receptor, ACE2, to more “open” intermediate conformations, with the S1 receptor binding domains (RBDs) in the “up” position (22,35–37,40,70). Proteolytic cleavage of the S gp in various contexts further activates the S gp, allowing it to achieve fusion-active conformations. The propensity of Class I viral envelope glycoproteins to make transitions from the pretriggered conformation to downstream conformations has been termed “reactivity” or “triggerability” (71). Studies of HIV-1 envelope glycoproteins have shown that high triggerability promotes efficient virus infection and broadened tropism, but can also render the virus more susceptible to antibody neutralization (72,73).

**FIG 13.**
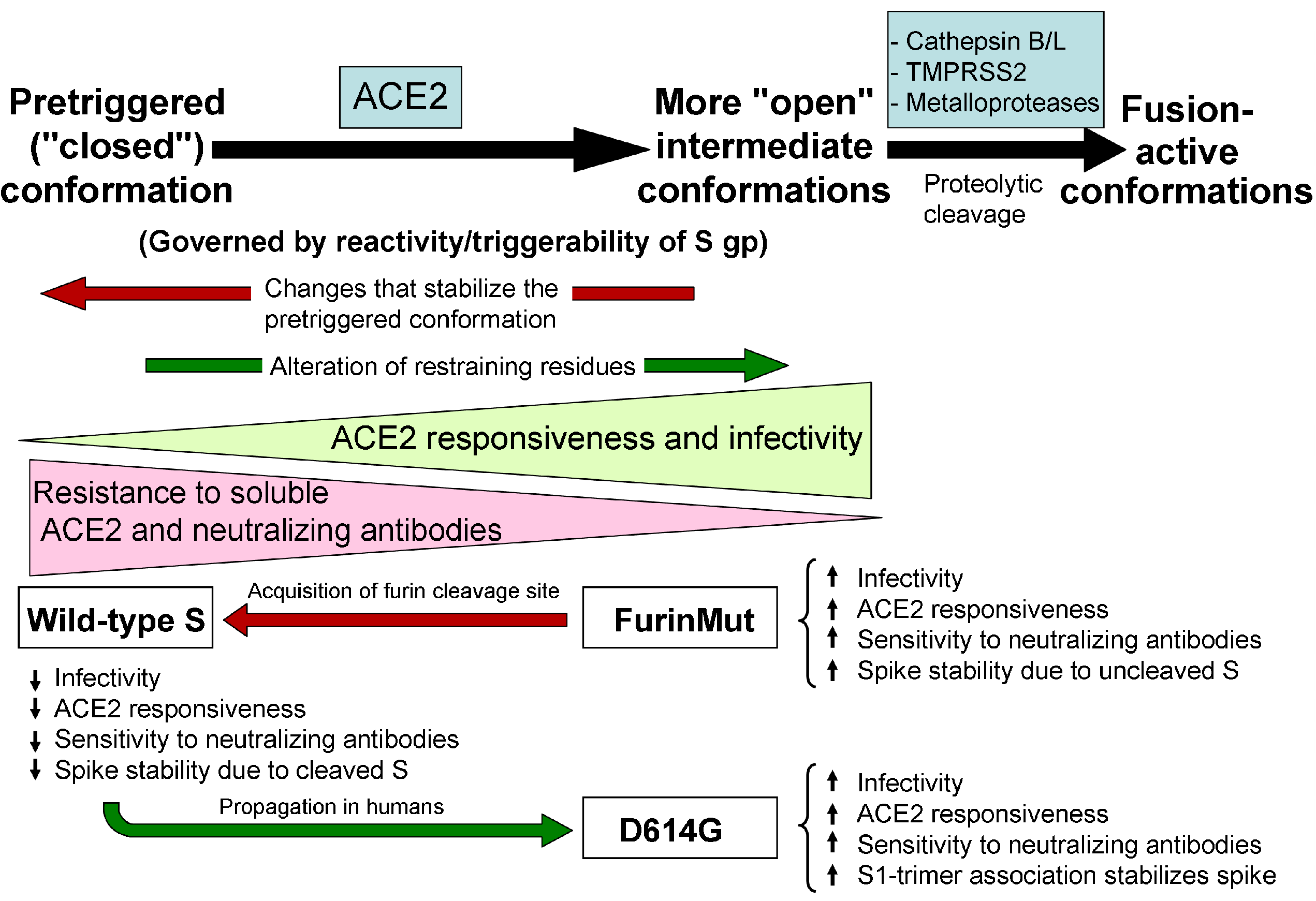
Model for the effects of two evolutionary changes on SARS-CoV-2 S gp function. The conformational transitions of the SARS-CoV-2 S gp during functional activation by ACE2 binding and proteolytic cleavage are depicted. The proposed effects of changes in S gp reactivity/triggerability on viral phenotypes follow from studies of other Class I viral envelope glycoproteins (33,72,73,93). The wild-type, FurinMut and D614G S glycoproteins are positioned along the pathway according to their phenotypes.

Our results indicate that two key features of the SARS-CoV-2 S gp, the furin cleavage site acquired during viral passage to humans from bat/intermediate hosts and the D614G change associated with increasing prevalence in human populations, influence related functions of the S gp. Acquisition of the furin cleavage site resulted in decreased infectivity and reduced sensitivity to soluble ACE2 and neutralizing antibodies, phenotypes suggestive of lowered triggerability (Fig. 13). Similarly, furin cleavage stabilizes the pretriggered conformation of the HIV-1 envelope glycoproteins (74–77). Cleavage of the wild-type SARS-CoV-2 S gp, unlike that of SARS-CoV-1, occurs in the S-expressing cell. Upon ACE2 binding, the wild-type SARS-CoV-2 S gp efficiently mediates syncytium formation and cytopathic effects, functions that are dramatically enhanced by S1-S2 cleavage. The furin cleavage site-dependent fusion of the membranes of infected cells with those of adjacent ACE2-expressing cells might also enhance cell-to-cell transmission via virus-cell synapses. Increased efficiency of cell-to-cell transmission could counterbalance the decreased cell-free infectivity of the viruses with the furin cleavage site. Increases in the efficiency of proteolytic processing of other Class I viral envelope glycoproteins have been associated with broadened viral tropism and enhanced virulence (78–86).

“Restraining residues” participate in molecular bonds that maintain Class I viral envelope glycoproteins in pretriggered conformations (72–73). Alteration of these restraining residues results in increased triggerability (Fig. 13). Asp 614 appears to fit the definition of a restraining residue. The D614G change increases virus infectivity and responsiveness to ACE2, but also results in moderate increases in sensitivity to neutralizing antisera. The proximity of Asp 614 (in the S1 C-terminal domain (CTD2)) to the S2 fusion peptide-proximal region (FPPR), which abuts the S1 CTD1 and has been proposed to suppress the opening of the S1 RBD (40), could account for these effects on viral phenotypes. Based on the reported structures of the SARS-CoV-2 spike glycoprotein (40,87), several amino acid residues (Lys 835, Gln 836 and Lys 854) are in close proximity to Asp 614 and therefore could potentially influence D614G phenotypes. However, alanine substitutions of these residues did not abolish the observed D614G phenotypes (data not shown). Of note, the D614G change partially compensates for the effects of acquisition of the furin cleavage site on virus infectivity and ACE2 responsiveness (Fig. 13). It will be of interest to evaluate whether other, less common evolutionary changes affect S gp conformational equilibrium (88).

The stability of functional viral spikes may significantly impact modes and efficiency of transmission. Based on the observation that soluble ACE2 induces shedding of the S1 glycoprotein from the SARS-CoV-2 S trimer, downstream “open” conformations may be more labile than the pretriggered conformation. The lability of open spike conformations could potentially nullify any replicative advantage of increased triggerability. The more triggerable FurinMut and D614G mutants solve this potential problem in different ways. By retaining a covalent bond between S1 and S2, FurinMut maintains and even increases spike stability. D614G retains an intact furin cleavage site but strengthens the association of S1 with the S trimer, possibly by improving S1 CTD2-S2 interactions. The resulting higher spike density allows FurinMut and D614G to take replicative advantage of increased ACE2 triggerability.

Proteolytic cleavage of the ACE2-bound S gp intermediate promotes conformational changes conducive to achieving membrane fusion. Depending on context, different proteases may be involved. During cell-free virus entry, cysteine proteases like Cathepsin B/L can act on endocytosed viruses. TMPRSS2 expressed on the surface of the ACE2-expressing target cells could enhance both cell-free virus infection and cell-cell fusion mediated by the SARS-CoV-2 S gp. TMPRSS2 enhancement of cell-cell fusion was particularly robust for S gp mutants that were not efficiently cleaved at the S1/S2 junction; that the S2′Mut was an exception hints that TMPRSS2 cleavage at the S2′ site may contribute to the observed enhancement. Our results implicate matrix metalloproteases in cell-cell fusion but not virus infection mediated by the SARS-CoV-2 S gp. Zinc metalloproteases have been implicated in neurovirulent murine coronavirus entry and cell-cell fusion (89). Cytopathic effects resulting from the formation of lethal syncytia may contribute to the damage of lung tissue during SARS-CoV-2 infection (90). As a number of matrix metalloproteases are expressed in pulmonary tissue and can be upregulated in response to inflammation (91), their potential involvement in SARS-CoV-2 pathogenesis should be further explored.

The sampling of downstream, open conformations by uncleaved Class I viral envelope glycoproteins can result in the elicitation of poorly neutralizing antibodies unable to recognize the native pretriggered conformation. In the case of HIV-1, uncleaved envelope glycoproteins can traffic to the cell surface by bypassing the conventional Golgi transport pathway, and therefore are efficiently presented to the host immune system (Zhang et al., unpublished observations). Likewise, substantial fractions of uncleaved SARS-CoV-2 S gp also appeared on the surface of overexpressing 293T cells, even when anterograde Golgi transport was blocked (Fig. 3D). The use of Golgi bypass pathways by conformationally flexible S glycoproteins could account for the observation that the vast majority of antibodies raised to the S gp during natural SARS-CoV-2 infection fail to neutralize the virus (27,92). Although limited to specific experimental systems, our study highlights the impact of SARS-CoV-2 S gp variation and host factors on spike synthesis, conformation and sensitivity to inhibition. These insights could assist the development of effective interventions.

## MATERIALS AND METHODS

### Cell lines

Human female embryonic kidney (HEK) 293T cells (ATCC) and COS-1 African green monkey male kidney fibroblasts (ATCC) were grown in Dulbecco modified Eagle medium (DMEM) supplemented with 10% fetal bovine serum (FBS) and 100 μg/ml penicillin-streptomycin (Pen-Strep) (Life Technologies). 293T cells were used for transient expression of the SARS-CoV-2 S gp variants, in some cases with HIV-1 Gag/protease or with SARS-CoV-2 M, E and N proteins. 293T cells transiently expressing omega-gal and either ACE2 or ACE2 + TMPRSS2 were used as target cells in the cell-cell fusion assay. COS-1 cells transiently expressing alpha-gal and SARS-CoV-2 S gp variants or HIV-1 envelope glycoproteins (AD8 or JR-FL strain) were used as effector cells in the cell-cell fusion assay. Cf2Th-CD4/CCR5 canine thymic epithelial cells expressing human CD4 and CCR5 coreceptors were used as target cells for evaluating cell-cell fusion mediated by the HIV-1 envelope glycoproteins; Cf2Th-CD4/CCR5 cells were grown in the above medium supplemented with 0.4 mg/ml G418 and 0.2 mg/ml hygromycin.

293T cells constitutively expressing human ACE2 (293T-ACE2 cells) were established by transducing 293T cells with the pLenti-III-hACE2 lentivirus vector (see below). The lentivirus vector was prepared by transfecting 1×10^6^ 293T cells in a six-well plate with μg psPAX2, 2 μg pLenti-III-hACE2 and 0.5 μg pVSVG using Lipofectamine 3000 (Thermo Fisher Scientific). Two days after transfection, the supernatant was harvested, spun at 5000 rpm for 15 minutes to remove cell debris, and then filtered (0.45-μm). The filtered supernatant containing recombinant lentiviral vectors was used to transduce 293T cells seeded one day before. Twenty-four hours later, the cells were selected in DMEM with 2 μg/ml puromycin with 1x Pen-Strep and 10% FBS for one week. A polyclonal puromycin-resistant population of 293T-ACE2 cells was shown to express human ACE2 by Western blotting. The 293T-ACE2 cells were used as target cells for infection by lentivirus and VSV pseudotypes, and as target cells in cell-cell fusion and syncytium formation assays.

The wild-type SARS-CoV-2 S gp, with Asp 614, was inducibly expressed in Lenti-x-293T human female kidney cells from Takara Bio (Catalog #: 632180). Lenti-x-293T cells were grown in DMEM with 10% heat-inactivated FBS supplemented with L-glutamine and Pen-Strep.

Lenti-x-293T cells constitutively expressing the reverse tetracycline-responsive transcriptional activator (rtTA) (Lenti-x-293T-rtTa cells (D1317)) (94) were used as the parental cells for the 293T-S and 293T-S-ACE2 cell lines. The 293T-S (D1481) cells inducibly expressing the wild-type SARS-CoV-2 S gp with a carboxy-terminal His_6_ tag were produced by transduction of Lenti-x-293T-rtTA cells with the K5648 recombinant lentivirus vector (see below). The packaged K5648 lentivirus vector (60 μl volume) was incubated with 2×10^5^ Lenti-x-293T-rtTA cells in DMEM, tumbling at 37°C overnight. The cells were then transferred to a 6-well plate in 3 ml DMEM/10% FBS/Pen-Strep and subsequently selected with 10 μg/ml puromycin.

293T-S-ACE2 (D1496) cells inducibly express the wild-type SARS-CoV-2 S gp and constitutively express human ACE2. Briefly, the 293T-S-ACE2 cells were produced by transduction of the 293T-S cells with the K5659 recombinant lentivirus vector, which expresses human ACE2 (see below). The packaged K5659 lentivirus vector (60 μl volume) was incubated with 2×10^5^ 293T-S cells in DMEM, tumbling at 37°C overnight. The cells were then transferred to a 6-well plate in 3 ml DMEM/10% FBS/Pen-Strep and subsequently selected with 10 μg/ml puromycin for four days.

### Plasmids

The wild-type and mutant SARS-CoV-2 S glycoproteins were expressed transiently by a pcDNA3.1(-) vector (Thermo Fisher Scientific). The wild-type SARS-CoV-2 spike (S) gene sequence, which encodes an aspartic acid residue at position 614, was obtained from the National Center for Biological Information (NC_045512.20). The gene was modified to encode a Gly_3_ linker and His_6_ tag at the carboxyl terminus. The modified S gene was codon-optimized, synthesized by Integrated DNA Technologies, and cloned into the pcDNA3.1(-) vector. S mutants were made using Q5 High-Fidelity 2X Master Mix and KLD Enzyme Mix for site-directed mutagenesis, according to the manufacturer’s protocol (New England Biolabs), and One-Shot TOP10 Competent Cells.

Inducible expression of the wild-type SARS-CoV-2 S gp was obtained using a self-inactivating lentivirus vector comprising TRE3g-SARS-CoV-2-Spike-6His.IRS6A.Puro-T2A-GFP (K5648) (95). Here, the expression of the codon-optimized, wild-type S gene is under the control of a tetracycline response element (TRE) promoter. The internal ribosome entry site (IRES) allows expression of puro.T2A.EGFP, in which puromycin N-acetyltransferase and enhanced green fluorescent protein (eGFP) are produced by self-cleavage at the Thosea asigna 2A (T2A) sequence.

Constitutive expression of human ACE2 in the 293T-S-ACE2 (D1496) cells was achieved using lentivirus vector (K5659) comprising hCMV-ACE2.IRES.puro. The ACE2 gene (obtained from Addgene [Catalog #: 1786]) was placed under the control of the human cytomegalovirus (hCMV) immediate early promoter. The vector also expresses puromycin N-acetyltransferase downstream of an IRES.

To package the recombinant K5648 and K5659 lentiviral vectors, 5×10^5^ Lenti-x-293T cells were transfected with 1.5 μg K5648 or K5659 vector/plasmid, 1.5 μg lentivirus packaging plasmid (expressing HIV-1 Gag, Pro, Pol, Tat and Rev), and 1 μg of a plasmid expressing the VSV G glycoprotein using the FuGene HD reagent (Promega). Forty-eight hours later, the cell supernatants were clarified by centrifugation at 3000 rpm and filtered (0.45-μm). Lentiviral vector particles were concentrated 25-fold by centrifugation at 100,000 x g for two hours. Packaged vector preparations were aliquoted in 60 μl volumes and stored at −80°C.

The pLenti-III-hACE2 plasmid was used to establish 293T-ACE2 cells constitutively expressing full-length human ACE2. A plasmid containing the ACE2 gene (Addgene) was digested with Nhe I and Kpn I and the fragment was ligated into the similarly digested pLenti-III-HA vector plasmid, producing pLenti-III-hACE2. To produce VSV G-pseudotyped lentiviruses encoding ACE2, 1×10^6^ 293T cells were transfected with 2 μg pLenti-III-hACE2, 1.5 μg psPAX2 and 0.5 μg pVSVG using Lipofectamine 3000 (Thermo Fisher Scientific). Two days later, the cell supernatant was harvested, clarified by centrifugation at 5000 rpm for 15 minutes, and filtered (0.45-μm). The recombinant lentiviruses in the filtered supernatants were used to transduce 293T cells, which were screened in DMEM/10% FBS/Pen-Strep with 2 μg/ml puromycin for one week. After selection of a polyclonal puromycin-resistant population, the expression of human ACE2 was confirmed by Western blotting.

The pcDNA3.1(-)-sACE2 plasmid expressing soluble ACE2 was made by Q5 site-directed mutagenesis (New England Biolabs) from the pcDNA3.1(-)-ACE2-Strep plasmid (Addgene, Catalog #: 1786), using the primers hACE2-strep-for: ccgcagtttgaaaaatagATATGGCTGATTGTTTTTGGAGTTG and hACE2-strep-rev: atggctccatcctcctccGGAAACAGGGGGCTGGTTAGGA

### Preparation of soluble ACE2

Expi293F cells, at a density of 3×10^6^ cells/ml, were transfected with 100 μg pcDNA3.1(-)-sACE2 using FectPRO DNA Transfection Reagent (PolyPlus-transfection). Three days later, cell supernatants were clarified by low-speed centrifugation (1,500 x g for 15 minutes), filtered (0.45-μm), and incubated with 1 ml Strep-Tactin resin (IBA Lifesciences) for 2 hours at 4°C with rotation. The mixture was applied to a Biorad column, washed with 30 bed volumes of 1x Strep-Tactin washing buffer and eluted with 10 bed volumes of 1x Strep-Tactin elution buffer. The eluate was concentrated using a 30-kD MWCO ultrafilter and then dialyzed twice against 1X PBS. The protein concentration was measured by the Bradford method (Thermo Fisher Scientific).

### Lentiviruses pseudotyped by S glycoproteins

Subconfluent 293T cells in a T75 flask were cotransfected with 1 μg of the S plasmid, 1 μg of the psPAX2 HIV-1 Gag-Pol packaging plasmid and 3 μg of the luciferase-expressing pLucX plasmid, using Effectene (Qiagen) according to the manufacturer’s protocol. Three to five days after transfection, cells and particles were processed.

### VSV pseudotyped by S glycoproteins

Subconfluent 293T cells in a T75 flask were transfected with 15 μg of the SARS-CoV-2 S plasmid using polyethylenimine (Polysciences), following the manufacturer’s protocol. Twenty-four hours later, cells were infected at a multiplicity of infection of 3-5 with rVSV-ΔG pseudovirus bearing a luciferase gene (Kerafast) for 2 hours at 37°C and then washed 6 times with 1X PBS. Cell supernatants containing S-pseudotyped VSV were harvested 24 hours later, clarified by low-speed centrifugation (2,000 rpm for 10 min), and either characterized immediately or stored at −80°C for later analysis. An additional 0.45-μm filtration step was included in some experiments to further purify the virus, but this step did not affect virus infectivity or neutralization patterns.

### S expression, processing and VLP incorporation

293T cells were transfected to produce lentivirus or VSV VLPs pseudotyped with S gp variants, as described above. At 3-5 days after transfection, a fraction of the cells was lysed and the cell lysates were analyzed by Western blotting, as described below. To evaluate the expression of S glycoproteins on the cell surface, cells were incubated with a 1:100 dilution of NYP01 convalescent serum for 1 hr at room temperature (RT). Cells were briefly spun to remove unbound antibody and lysed. Clarified lysates were incubated with Protein A-agarose beads for 1 hr at room temperature. The beads were washed three times and boiled, after which the captured proteins were analyzed by Western blot. To prepare VLPs, cell supernatants were collected, filtered (0.45-μm), and pelleted at 100,000 x g (for lentivirus VLPs) or 14,000 x g (for VSV VLPs) for 1 hr at 4°C. Samples were Western blotted with 1:2,000 dilutions of rabbit anti-SARS-Spike S1, mouse anti-SARS-Spike S1, rabbit anti-SARS-Spike S2, rabbit anti-p55/p24/p17 or mouse anti-VSV-NP or a 1:10,000 dilution of mouse anti-β-actin as the primary antibodies. HRP-conjugated anti-rabbit or anti-mouse antibodies at a slightly lower dilution were used as secondary antibodies in the Western blots. The adjusted integrated volumes of S, S1 and S2 bands from unsaturated Western blots were calculated using Bio-Rad Image Lab Software. The values for the expression of mutant S glycoproteins were calculated and normalized to the values for the wild-type S gp (WT) as follows:

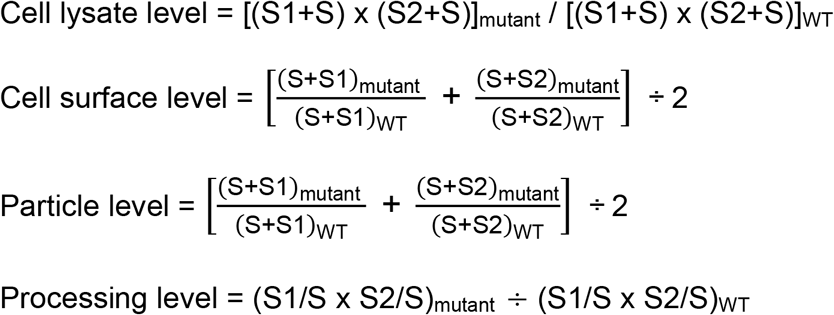

### Deglycosylation of S glycoproteins

S glycoproteins in cell lysates or on cell surfaces or VLPs were prepared as described above. Protein samples were boiled in 1X Denaturing Buffer and incubated with PNGase F or Endo Hf (New England Biolabs) for 1-1.5 hr at 37°C following the manufacturer’s protocol. The samples were then analyzed by SDS-PAGE and Western blotting.

### Effects of SARS-CoV-2 M/E/N proteins

293T cells were cotransfected with SARS-CoV-2 S, M, E and N plasmids at equimolar concentrations. In the mock sample, an equivalent amount of the pcDNA3.1(-) plasmid was used in place of the M,E and N plasmids. In experiments using Brefeldin A (BFA), BFA was added at 6 hr after transfection. At 3 days after transfection, samples from cell lysates and on cell surfaces and VLPs were prepared as described above.

### S1 shedding from S gp-expressing cells

293T cells were transfected with pcDNA3.1(-) plasmids expressing the wild-type and mutant SARS-CoV-2 S glycoproteins, using Effectene according to the manufacturer’s protocol. Cell supernatants were collected, filtered (0.45-μm) and incubated with a 1:100 dilution of NYP01 convalescent serum and Protein A-agarose beads for 1-2 hr at room temperature. Beads were washed three times and samples were Western blotted with a mouse anti-S1 antibody. Band intensity was determined as described above. The subunit association index of each mutant was calculated as follows:

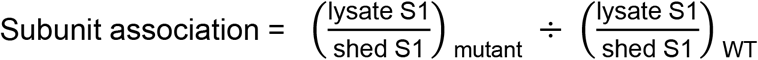

### Cell cytotoxicity/MTT assay

293T-S cells and 293T-S-ACE2 cells seeded in clear 96-well plates at different densities were induced with 1 μg/ml doxycycline. Cell viability was measured using the MTT assay (Abcam) at different times following induction, according to the manufacturer’s protocol.

### Syncytium formation and inhibition

293T-ACE2 cells were cotransfected with an eGFP-expressing plasmid and either pcDNA3.1(-) or a pcDNA3.1 plasmid expressing the wild-type or mutant SARS-CoV-2 S gp. For the HIV-1 control, 293T cells were transfected with plasmids expressing eGFP, the HIV-1_JR-FL_ envelope glycoproteins, CD4 and CCR5. Twenty-four hours after transfection, cells were imaged using a fluorescence microscope with a green light filter. In the syncytium inhibition assay, 10 μM ilomastat, 10 μM marimastat or 100 μM camostat mesylate was added during transfection and cells were imaged at different time points after transfection.

### Cell-cell fusion (alpha-complementation) assay

COS-1 effector cells plated in black-and-white 96-well plates were cotransfected with a plasmid expressing alpha-gal and either pcDNA3.1 or plasmids expressing the SARS-CoV-1 S gp or wild-type or mutant SARS-CoV-2 S glycoproteins at a 1:1 ratio, using Effectene according to the manufacturer’s protocol. At the same time, 293T target cells plated in 6-well plates were cotransfected with plasmids expressing omega-gal and ACE2 at a 1:1 ratio, using Effectene. In the experiments shown in Fig. 12A and B, 293T target cells were cotransfected with equimolar concentrations of plasmids expressing omega-gal, ACE2 and TMPRSS2. Forty-eight hours after transfection, target cells were scraped and resuspended in medium. Medium was removed from the effector cells, and target cells were then added to effector cells (one target-cell well provides sufficient cells for 50 effector-cell wells). Plates were spun at 500 x g for 3 minutes and then incubated at 37 °C in 5% CO_2_ for 3-4 hours. Medium was aspirated and cells were lysed in Tropix lysis buffer (Thermo Fisher Scientific). The β-galactosidase activity in the cell lysates was measured using the Galacto-Star Reaction Buffer Diluent with Galacto-Star Substrate (Thermo Fisher Scientific) following the manufacturer’s protocol.

In the inhibition assay, to allow for optimal contact, target cells were scraped and resuspended in medium, and inhibitors were added at the indicated concentrations. Cells were then incubated at 37 °C in 5% CO_2_ for 2 hr before they were added to effector cells, as described above.

For the HIV-1 envelope glycoprotein control, effector cells were cotransfected with plasmids expressing alpha-gal and the HIV-1_AD8_ Env and Tat at a 1:1:0.125 ratio. 293T target cells were cotransfected with plasmids expressing omega-gal, CD4 and CCR5 at a 1:1:1 ratio.

A variation of the above protocol was used for the cell-cell fusion assays shown in Fig. 12B and D. For the experiment shown in Fig. 12B, 293T-S effector cells in black-and-white 96-well plates were transfected with a plasmid expressing alpha-gal using Lipofectamine 3000 and either induced with 1 μg/ml doxycycline or mock treated. In parallel, 293T-ACE2 target cells in 6-well plates were cotransfected with a plasmid expressing omega-gal and a plasmid expressing either TMPRSS2 or, as a control, pcDNA3.1(-). One day after transfection, the target cells were treated with 5 mM EDTA in 1X PBS, resuspended in fresh medium, and aliquoted. The target cells were either untreated as a control or incubated with camostat mesylate, marimastat or a combination of camostat and marimastat, and then added to effector cells. For the experiment shown in Fig. 12D, COS-1 cells were used as effector cells. The effector cells were transfected with a plasmid expressing alpha-gal and a plasmid expressing the HIV-1_JR-FL_ envelope glycoprotein or the wild-type SARS-CoV-2 S gp, or pcDNA3.1 as a control. For the SARS-CoV-2 S gp-expressing effector cells, 293T-ACE2 cells were used as target cells; for effector cells expressing the HIV-1_JR-FL_ envelope glycoprotein, Cf2Th-CD4/CCR5 cells were used as target cells. The target cells were transfected with a plasmid expressing omega-gal. Alpha-complementation assays were performed one day after transfection, as described above.

### Virus infectivity and cold sensitivity

Luciferase-containing recombinant lentivirus and VSV-ΔG vectors pseudotyped with SARS-CoV-2 S gp variants were produced as described above. The recombinant viruses were incubated with 293T-ACE2 cells, and 24-48 hours later, luciferase activity in the cells was measured. To measure cold sensitivity, the viruses were incubated on ice for various lengths of time prior to measuring their infectivity, as described above.

### Virus neutralization by sACE2 and sera

Neutralization assays were performed by adding 200-300 TCID_50_ of rVSV-ΔG pseudotyped with SARS-CoV-2 S gp variants into serial dilutions of sACE2 and sera. The mixture was dispensed onto a 96-well plate in triplicate and incubated for 1 h at 37°C. Approximately 7 × 10^4^ 293T-ACE2 cells were then added to each well, and the cultures were maintained for an additional 24 h at 37°C before luciferase activity was measured. Neutralization activity was calculated from the reduction in luciferase activity compared to controls. The concentrations of sACE2 and dilutions of sera that inhibited 50% of infection (the IC_50_ and ID_50_ values, respectively) were determined by fitting the data in five-parameter dose-response curves using GraphPad Prism 8 (GraphPad Software Inc.).

### Stability of spike protein on VLPs

To prepare lentivirus particles containing wild-type or D614G S gp, ~7×10^6^ 293T cells in T75 flasks were transfected with 7.5 μg psPAX2 and 7.5 μg of the S-expressing pcDNA3.1 plasmid using Lipofectamine 3000. Mock controls did not include the psPAX2 plasmid. Two days later, cell supernatants were collected, clarified by centrifugation at 1,500 x g for 15 min and filtered (0.45-μm). VLP-containing supernatants were incubated at 37°C for various lengths of time. Then the VLPs were pelleted at 14,000 x g for 1 hr at 4°C. The pellets were suspended in 50 μl of 1X LDS buffer containing 100 mM DTT, and Western blotted using rabbit anti-S1, rabbit anti-S2 and anti-p55/p24/p17 antibodies.

### Interaction of soluble ACE2 with S on VLPs

VLPs containing the wild-type or D614G S gp prepared as described above were used to measure the binding of soluble ACE2 (sACE2) to the viral spike, and to study the effect of sACE2 binding on the shedding of the S1 gp from the spike. After clarification and filtration, the VLP-containing cell supernatants were incubated with different concentrations of sACE2 at either 0°C or 37°C for one hour. Afterwards, the VLPs were pelleted at 14,000 x g for 1 hr at 4°C, and analyzed by Western blotting as described above.

### S1 association with solubilized S gp trimers

293T cells were transfected with pcDNA3.1 plasmids expressing the wild-type or D614G SARS-CoV-2 S glycoproteins, which contain carboxy-terminal His_6_ tags, using Lipofectamine 3000. Two days after transfection, cells were washed and lysed in lysis buffer containing 1% Cymal-5 on ice for 5 minutes. Clarified cell lysates were aliquoted and incubated at different temperatures for 1 hr, cooled on ice and then incubated with Ni-NTA Superflow beads (Thermo Fisher Scientific) for 1 hr at 4°C. Beads were washed 3 times with lysis buffer at 4°C. After washing, beads were resuspended in 1X LDS and 100 mM DTT, boiled for 5 minutes and Western blotted with anti-S1 and anti-S2 antibodies.

### Palmitoylation assay

Potential palmitoylation sites on the SARS-CoV-2 S2 glycoprotein were studied using acyl-PEG exchange (68). Briefly, 293T cells were transfected with pcDNA3.1 plasmids expressing the SARS-CoV-2 S gp variants. Cell lysates in 4% SDS (~100 μg total protein) were treated with 10 mM Tris(2-carboxyethyl) phosphine hydrochloride and rotated at 350 rpm for 30 min at room temperature. Then, 25 mM NEM was added and the sample was incubated at room temperature for 2 hr. The reaction was terminated and washed with prechilled methanol-chloroform-H_2_O (4:1.5:3) sequentially three times. The protein precipitates were dissolved in 60 μl of TEA buffer with 4 mM EDTA and 4% SDS. Half of the dissolved protein solution was mixed with 1 M hydroxylamine in TEA buffer (pH 7.3) + 0.2% Triton X-100, while the remaining half was mixed with TEA buffer + 0.2% Triton X-100 followed by a 350-rpm rotation for 1 hr at room temperature. Reactions were stopped by adding prechilled methanol-chloroform-H_2_O as mentioned above, and the precipitated pellets were dissolved in TEA buffer + 4 mM EDTA + 4% SDS by incubation at 50°C for 10 minutes. The protein solutions were mixed with 1.33 mM mPEG-mal in TEA + 0.2% Triton X-100 and incubated at 25°C for 2 hr by nutation at 350 rpm. Reactions were stopped by adding prechilled methanol-chloroform-H_2_O as mentioned above. The protein pellets were dissolved with 1X LDS buffer + 100 mM DTT and subjected to Western blot analysis using rabbit anti-S2 antibody.

### Statistical analysis

Two-tailed Student’s t-tests were used to determine statistical significance. Significant differences are indicated in the figures as follows (*, P < 0.05; **, P < 0.01; ***, P < 0.001).

## ACKNOWLEDGMENTS

We thank Ms. Elizabeth Carpelan for manuscript preparation. We thank Drs. Lihong Liu and David Ho (Columbia University) and Peihui Wang (Shandong University) for reagents. This work was supported by the Genetic Sequencing Core of the University of Alabama at Birmingham (UAB) Center for AIDS Research (NIH P30 AI27767), grants from the National Institutes of Health (AI145547 and AI124982) and by a gift from the late William F. McCarty-Cooper.

## REFERENCES

1. Cui, J., Li, F., and Shi, Z.L. (2019). Origin and evolution of pathogenic coronaviruses. Nat Rev Microbiol 17, 181–192.

2. de Wit, E., van Doremalen, N., Falzarano, D., and Munster, V.J. (2016). SARS and MERS: recent insights into emerging coronaviruses. Nat Rev Microbiol 14, 523–534.

3. Graham, R.L., and Baric, R.S. (2010). Recombination, reservoirs, and the modular spike: mechanisms of coronavirus cross-species transmission. J Virol 84, 3134–3146.

4. Perlman, S., and Netland, J. (2009). Coronaviruses post-SARS: update on replication and pathogenesis. Nat Rev Microbiol 7, 439–450.

5. Song, Z., Xu, Y., Bao, L., Zhang, L., Yu, P., Qu, Y., Zhu, H., Zhao, W., Han, Y., and Qin, C. (2019). From SARS to MERS, thrusting coronaviruses into the spotlight. Viruses. 2019 Jan 14;11(1):59.

6. Li, Q., Guan, X., Wu, P., Wang, X., Zhou, L., Tong, Y., Ren, R., Leung, K.S.M., Lau, E.H.Y., Wong, J.Y., et al. (2020). Early transmission dynamics in Wuhan, China, of novel coronavirus-infected pneumonia. N Engl J Med 382, 1199–1207.

7. Huang, C., Wang, Y., Li, X., Ren, L., Zhao, J., Hu, Y., Zhang, L., Fan, G., Xu, J., Gu, X., et al. (2020). Clinical features of patients infected with 2019 novel coronavirus in Wuhan, China. Lancet 395, 497–506.

8. Wu, F., Zhao, S., Yu, B., Chen, Y.M., Wang, W., Song, Z.G., Hu, Y., Tao, Z.W., Tian, J.H., Pei, Y.Y., et al. (2020). A new coronavirus associated with human respiratory disease in China. Nature 579, 265–269.

9. Zhou, P., Yang, X.L., Wang, X.G., Hu, B., Zhang, L., Zhang, W., Si, H.R., Zhu, Y., Li, B., Huang, C.L., et al. (2020). A pneumonia outbreak associated with a new coronavirus of probable bat origin. Nature 579, 270–273.

10. Coronaviridae Study Group of the International Committee on Taxonomy of, V. (2020). The species severe acute respiratory syndrome-related coronavirus: classifying 2019-nCoV and naming it SARS-CoV-2. Nat Microbiol 5, 536–544.

11. Lv, M., Luo, X., Estill, J., Liu, Y., Ren, M., Wang, J., Wang, Q., Zhao, S., Wang, X., Yang, S., et al. (2020). Coronavirus disease (COVID-19): a scoping review. Euro Surveill 25, 2000125.

12. Dowd, J.B., Andriano, L., Brazel, D.M., Rotondi, V., Block, P., Ding, X., Liu, Y., and Mills, M.C. (2020). Demographic science aids in understanding the spread and fatality rates of COVID-19. Proc Natl Acad Sci U S A 117, 9696–9698.

13. Woo, P.C., Lau, S.K., Lam, C.S., Lau, C.C., Tsang, A.K., Lau, J.H., Bai, R., Teng, J.L., Tsang, C.C., Wang, M., et al. (2012). Discovery of seven novel mammalian and avian coronaviruses in the genus deltacoronavirus supports bat coronaviruses as the gene source of alphacoronavirus and betacoronavirus and avian coronaviruses as the gene source of gammacoronavirus and deltacoronavirus. J Virol 86, 3995–4008.

14. Huynh, J., Li, S., Yount, B., Smith, A., Sturges, L., Olsen, J.C., Nagel, J., Johnson, J.B., Agnihothram, S., Gates, J.E., et al. (2012). Evidence supporting a zoonotic origin of human coronavirus strain NL63. J Virol 86, 12816–12825.

15. Jaimes, J.A., Andre, N.M., Chappie, J.S., Millet, J.K., and Whittaker, G.R. (2020). Phylogenetic analysis and structural modeling of SARS-CoV-2 spike protein reveals an evolutionary distinct and proteolytically sensitive activation loop. J Mol Biol 432, 3309–3325.

16. Kissler, S.M., Tedijanto, C., Goldstein, E., Grad, Y.H., and Lipsitch, M. (2020). Projecting the transmission dynamics of SARS-CoV-2 through the postpandemic period. Science 368, 860–868.

17. Venkat Kumar, G., Jeyanthi, V., and Ramakrishnan, S. (2020). A short review on antibody therapy for COVID-19. New Microbes New Infect 35, 100682.

18. Wang, C., Li, W., Drabek, D., Okba, N.M.A., van Haperen, R., Osterhaus, A., van Kuppeveld, F.J.M., Haagmans, B.L., Grosveld, F., and Bosch, B.J. (2020). A human monoclonal antibody blocking SARS-CoV-2 infection. Nat Commun 11, 2251.

19. Callaway, E. (2020). The race for coronavirus vaccines: a graphical guide. Nature 580, 576–577.

20. Billington, J., Deschamps, I., Erck, S.C., Gerberding, J.L., Hanon, E., Ivol, S., Shiver, J.W., Spencer, J.A., and Van Hoof, J. (2020). Developing vaccines for SARS-CoV-2 and future epidemics and pandemics: applying lessons from past outbreaks. Health Secur 18, 241–249.

21. Padron-Regalado, E. (2020). Vaccines for SARS-CoV-2: lessons from other coronavirus strains. Infect Dis Ther 9, 1–20.

22. Walls, A.C., Park, Y.J., Tortorici, M.A., Wall, A., McGuire, A.T., and Veesler, D. (2020). Structure, function, and antigenicity of the SARS-CoV-2 spike glycoprotein. Cell 181, 281–292 e286.

23. Yuan, M., Wu, N.C., Zhu, X., Lee, C.D., So, R.T.Y., Lv, H., Mok, C.K.P., and Wilson, I.A. (2020). A highly conserved cryptic epitope in the receptor binding domains of SARS-CoV-2 and SARS-CoV. Science 368, 630–633.

24. Quinlan, B.D., Mou, H., Zhang, L., Guo, Y., He, W., Ojha, A., Parcells, M.S., Luo, G., Li, W., Zhong, G., et al. (2020). The SARS-CoV-2 receptor-binding domain elicits a potent neutralizing response without antibody-dependent enhancement. bioRxiv. doi: 10.1101/2020.04.10.036418.

25. Robbiani, D.F., Gaebler, C., Muecksch, F., Lorenzi, J.C.C., Wang, Z., Cho, A., Agudelo, M., Barnes, C.O., Gazumyan, A., Finkin, S., et al. (2020). Convergent antibody responses to SARS-CoV-2 in convalescent individuals. Nature 584, 437–442.

26. Rogers, T.F., Zhao, F., Huang, D., Beutler, N., Burns, A., He, W.T., Limbo, O., Smith, C., Song, G., Woehl, J., et al. (2020). Isolation of potent SARS-CoV-2 neutralizing antibodies and protection from disease in a small animal model. Science 369, 956–963.

27. Wec, A.Z., Wrapp, D., Herbert, A.S., Maurer, D.P., Haslwanter, D., Sakharkar, M., Jangra, R.K., Dieterle, M.E., Lilov, A., Huang, D., et al. (2020). Broad neutralization of SARS-related viruses by human monoclonal antibodies. Science 369, 731–736.

28. Zost, S.J., Gilchuk, P., Case, J.B., Binshtein, E., Chen, R.E., Nkolola, J.P., Schafer, A., Reidy, J.X., Trivette, A., Nargi, R.S., et al. (2020). Potently neutralizing and protective human antibodies against SARS-CoV-2. Nature 584, 443–449.

29. Liu, L., Wang, P., Nair, M.S., Yu, J., Rapp, M., Wang, Q., Luo, Y., Chan, J.F., Sahi, V., Figueroa, A., et al. (2020). Potent neutralizing antibodies against multiple epitopes on SARS-CoV-2 spike. Nature 584, 450–456.

30. Yu, J., Tostanoski, L.H., Peter, L., Mercado, N.B., McMahan, K., Mahrokhian, S.H., Nkolola, J.P., Liu, J., Li, Z., Chandrashekar, A., et al. (2020). DNA vaccine protection against SARS-CoV-2 in rhesus macaques. Science 369, 806–811.

31. Mercado, N.B., Zahn, R., Wegmann, F., Loos, C., Chandrashekar, A., Yu, J., Liu, J., Peter, L., McMahan, K., Tostanoski, L.H., et al. (2020). Single-shot Ad26 vaccine protects against SARS-CoV-2 in rhesus macaques. Nature (2020). https://doi.org/10.1038/s41586-020-2607-z

32. Ou, X., Liu, Y., Lei, X., Li, P., Mi, D., Ren, L., Guo, L., Guo, R., Chen, T., Hu, J., et al. (2020). Characterization of spike glycoprotein of SARS-CoV-2 on virus entry and its immune cross-reactivity with SARS-CoV. Nat Commun 11, 1620 (2020).

33. Tortorici, M.A., and Veesler, D. (2019). Structural insights into coronavirus entry. Adv Virus Res 105, 93–116.

34. Li, F. (2016). Structure, function, and evolution of coronavirus spike proteins. Annu Rev Virol 3, 237–261.

35. Wrapp, D., Wang, N., Corbett, K.S., Goldsmith, J.A., Hsieh, C.L., Abiona, O., Graham, B.S., and McLellan, J.S. (2020). Cryo-EM structure of the 2019-nCoV spike in the prefusion conformation. Science 367, 1260–1263.

36. Shang, J., Ye, G., Shi, K., Wan, Y., Luo, C., Aihara, H., Geng, Q., Auerbach, A., and Li, F. (2020). Structural basis of receptor recognition by SARS-CoV-2. Nature 581, 221–224.

37. Yan, R., Zhang, Y., Li, Y., Xia, L., Guo, Y., and Zhou, Q. (2020). Structural basis for the recognition of SARS-CoV-2 by full-length human ACE2. Science 367, 1444–1448.

38. Hoffmann, M., Kleine-Weber, H., Schroeder, S., Kruger, N., Herrler, T., Erichsen, S., Schiergens, T.S., Herrler, G., Wu, N.H., Nitsche, A., et al. (2020). SARS-CoV-2 cell entry depends on ACE2 and TMPRSS2 and is blocked by a clinically proven protease inhibitor. Cell 181, 271–280 e278.

39. Xia, S., Liu, M., Wang, C., Xu, W., Lan, Q., Feng, S., Qi, F., Bao, L., Du, L., Liu, S., et al. (2020). Inhibition of SARS-CoV-2 (previously 2019-nCoV) infection by a highly potent pan-coronavirus fusion inhibitor targeting its spike protein that harbors a high capacity to mediate membrane fusion. Cell Res 30, 343–355.

40. Cai, Y., Zhang, J., Xiao, T., Peng, H., Sterling, S.M., Walsh, R.M., Rawson, S., Rits-Volloch, S., and Chen, B. (2020). Distinct conformational states of SARS-CoV-2 spike protein. Science 25 Sep 2020: Vol. 369, Issue 6511, pp. 1586–1592. DOI: 10.1126/science.abd4251.

41. Coutard, B., Valle, C., de Lamballerie, X., Canard, B., Seidah, N.G., and Decroly, E. (2020). The spike glycoprotein of the new coronavirus 2019-nCoV contains a furin-like cleavage site absent in CoV of the same clade. Antiviral Res 176, 104742.

42. Hoffmann, M., Kleine-Weber, H., and Pohlmann, S. (2020). A multibasic cleavage site in the spike protein of SARS-CoV-2 is essential for infection of human lung cells. Mol Cell 78, 779–784 e775.

43. Korber, B., Fischer, W.M., Gnanakaran, S., Yoon, H., Theiler, J., Abfalterer, W., Hengartner, N., Giorgi, E.E., Bhattacharya, T., Foley, B., et al. (2020). Tracking changes in SARS-CoV-2 spike: evidence that D614G increases infectivity of the COVID-19 virus. Cell 182, 812–827 e819.

44. Zhang, L., Jackson, C.B., Mou, H., Ojha, A., Rangarajan, E.S., Izard, T., Farzan, M., and Choe, H. (2020). The D614G mutation in the SARS-CoV-2 spike protein reduces S1 shedding and increases infectivity. bioRxiv. doi: 10.1101/2020.06.12.148726.

45. Lontok, E., Corse, E., and Machamer, C.E. (2004). Intracellular targeting signals contribute to localization of coronavirus spike proteins near the virus assembly site. J Virol 78, 5913–5922.

46. McBride, C.E., Li, J., and Machamer, C.E. (2007). The cytoplasmic tail of the severe acute respiratory syndrome coronavirus spike protein contains a novel endoplasmic reticulum retrieval signal that binds COPI and promotes interaction with membrane protein. J Virol 81, 2418–2428.

47. Stertz, S., Reichelt, M., Spiegel, M., Kuri, T., Martinez-Sobrido, L., Garcia-Sastre, A., Weber, F., and Kochs, G. (2007). The intracellular sites of early replication and budding of SARS-coronavirus. Virology 361, 304–315.

48. Ujike, M., Huang, C., Shirato, K., Makino, S., and Taguchi, F. (2016). The contribution of the cytoplasmic retrieval signal of severe acute respiratory syndrome coronavirus to intracellular accumulation of S proteins and incorporation of S protein into virus-like particles. J Gen Virol 97, 1853–1864.

49. Petit, C.M., Chouljenko, V.N., Iyer, A., Colgrove, R., Farzan, M., Knipe, D.M., and Kousoulas, K.G. (2007). Palmitoylation of the cysteine-rich endodomain of the SARS-coronavirus spike glycoprotein is important for spike-mediated cell fusion. Virology 360, 264–274.

50. McBride, C.E., and Machamer, C.E. (2010). Palmitoylation of SARS-CoV S protein is necessary for partitioning into detergent-resistant membranes and cell-cell fusion but not interaction with M protein. Virology 405, 139–148.

51. Stanley, P. (2011). Golgi glycosylation. Cold Spring Harb Perspect Biol 3, a005199.

52. Masters, P.S. (2006). The molecular biology of coronaviruses. Adv Virus Res 66, 193–292.

53. Schoeman, D., and Fielding, B.C. (2019). Coronavirus envelope protein: current knowledge. Virol J 16, 69.

54. Nieto-Torres, J.L., Dediego, M.L., Alvarez, E., Jimenez-Guardeno, J.M., Regla-Nava, J.A., Llorente, M., Kremer, L., Shuo, S., and Enjuanes, L. (2011). Subcellular location and topology of severe acute respiratory syndrome coronavirus envelope protein. Virology 415, 69–82.

55. Mortola, E., and Roy, P. (2004). Efficient assembly and release of SARS coronavirus-like particles by a heterologous expression system. FEBS Lett 576, 174–178.

56. Siu, Y.L., Teoh, K.T., Lo, J., Chan, C.M., Kien, F., Escriou, N., Tsao, S.W., Nicholls, J.M., Altmeyer, R., Peiris, J.S., et al. (2008). The M, E, and N structural proteins of the severe acute respiratory syndrome coronavirus are required for efficient assembly, trafficking, and release of virus-like particles. J Virol 82, 11318–11330.

57. Helms, J.B., and Rothman, J.E. (1992). Inhibition by brefeldin A of a Golgi membrane enzyme that catalyses exchange of guanine nucleotide bound to ARF. Nature 360, 352–354.

58. Lippincott-Schwartz, J., Yuan, L.C., Bonifacino, J.S., and Klausner, R.D. (1989). Rapid redistribution of Golgi proteins into the ER in cells treated with brefeldin A: evidence for membrane cycling from Golgi to ER. Cell 56, 801–813.

59. Lippincott-Schwartz, J., Yuan, L., Tipper, C., Amherdt, M., Orci, L., and Klausner, R.D. (1991). Brefeldin A’s effects on endosomes, lysosomes, and the TGN suggest a general mechanism for regulating organelle structure and membrane traffic. Cell 67, 601–616.

60. Holland, A.U., Munk, C., Lucero, G.R., Nguyen, L.D., and Landau, N.R. (2004). Alpha-complementation assay for HIV envelope glycoprotein-mediated fusion. Virology 319, 343–352.

61. Johnson, M.C, Lyddon, T.D., Suarez, R., Salcedo, B., LePique, M., Graham, M., Ricana, C., Robinson, C., Ritter, D.G. (2020). Optimized Pseudotyping Conditions for the SARS-COV-2 Spike Glycoprotein. DOI: 10.1128/JVI.01062-20.

62. Bosch, B.J., Bartelink, W., and Rottier, P.J. (2008). Cathepsin L functionally cleaves the severe acute respiratory syndrome coronavirus class I fusion protein upstream of rather than adjacent to the fusion peptide. J Virol 82, 8887–8890.

63. Matsuyama, S., Ujike, M., Morikawa, S., Tashiro, M., and Taguchi, F. (2005). Protease-mediated enhancement of severe acute respiratory syndrome coronavirus infection. Proc Natl Acad Sci U S A 102, 12543–12547.

64. Millet, J.K., and Whittaker, G.R. (2015). Host cell proteases: critical determinants of coronavirus tropism and pathogenesis. Virus Res 202, 120–134.

65. Sainz, B., Jr., Rausch, J.M., Gallaher, W.R., Garry, R.F., and Wimley, W.C. (2005). Identification and characterization of the putative fusion peptide of the severe acute respiratory syndrome-associated coronavirus spike protein. J Virol 79, 7195–7206.

66. Madu, I.G., Roth, S.L., Belouzard, S., and Whittaker, G.R. (2009). Characterization of a highly conserved domain within the severe acute respiratory syndrome coronavirus spike protein S2 domain with characteristics of a viral fusion peptide. J Virol 83, 7411–7421.

67. Ou, X., Zheng, W., Shan, Y., Mu, Z., Dominguez, S.R., Holmes, K.V., and Qian, Z. (2016). Identification of the fusion peptide-containing region in betacoronavirus spike glycoproteins. J Virol 90, 5586–5600.

68. Percher, A., Ramakrishnan, S., Thinon, E., Yuan, X., Yount, J.S., and Hang, H.C. (2016). Mass-tag labeling reveals site-specific and endogenous levels of protein S-fatty acylation. Proc Natl Acad Sci U S A 113, 4302–4307.

69. Sergeeva, O.A., van der Goot, F.G. (2019). Anthrax toxin requires ZDHHC5-mediated palmitoylation of its surface-processing host enzymes. PNAS Jan 2019, 116 (4) 1279–1288.

70. Ke, Z., Oton, J., Qu, K., Cortese, M., Zila, V., McKeane, L., Nakane, T., Zivanov, J., Neufeldt, C.J., Cerikan, B., et al. (2020). Structures and distributions of SARS-CoV-2 spike proteins on intact virions. Nature. doi: 10.1038/s41586-020-2665-2.

71. Haim, H., Strack, B., Kassa, A., Madani, N., Wang, L., Courter, J.R., Princiotto, A., McGee, K., Pacheco, B., Seaman, M.S., et al. (2011). Contribution of intrinsic reactivity of the HIV-1 envelope glycoproteins to CD4-independent infection and global inhibitor sensitivity. PLoS Pathog 7, e1002101.

72. Herschhorn, A., Ma, X., Gu, C., Ventura, J.D., Castillo-Menendez, L., Melillo, B., Terry, D.S., Smith, A.B. III, Blanchard, S.C., Munro, J.B., et al. (2016). Release of gp120 restraints leads to an entry-competent intermediate state of the HIV-1 envelope glycoproteins. mBio 7, e01598–16.

73. Herschhorn, A., Gu, C., Moraca, F., Ma, X., Farrell, M., Smith, A.B. III, Pancera, M., Kwong, P.D., Schon, A., Freire, E., et al. (2017). The beta20-beta21 of gp120 is a regulatory switch for HIV-1 Env conformational transitions. Nat Commun 8, 1049.

74. Castillo-Menendez, L.R., Witt, K., Espy, N., Princiotto, A., Madani, N., Pacheco, B., Finzi, A., and Sodroski, J. (2018). Comparison of uncleaved and mature human immunodeficiency virus membrane envelope glycoprotein trimers. J Virol 92, e00277–18.

75. Haim, H., Salas, I., and Sodroski, J. (2013). Proteolytic processing of the human immunodeficiency virus envelope glycoprotein precursor decreases conformational flexibility. J Virol 87, 1884–1889.

76. Lu, M., Ma, X., Reichard, N., Terry, D.S., Arthos, J., Smith, A.B. III, Sodroski, J.G., Blanchard, S.C., and Mothes, W. (2020). Shedding-resistant HIV-1 envelope glycoproteins adopt downstream conformations that remain responsive to conformation-preferring ligands. J Virol 94, e00597–20.

77. Pancera, M., and Wyatt, R. (2005). Selective recognition of oligomeric HIV-1 primary isolate envelope glycoproteins by potently neutralizing ligands requires efficient precursor cleavage. Virology 332, 145–156.

78. Andre, N.M., Cossic, B., Davies, E., Miller, A.D., and Whittaker, G.R. (2019). Distinct mutation in the feline coronavirus spike protein cleavage activation site in a cat with feline infectious peritonitis-associated meningoencephalomyelitis. JFMS Open Rep 5, 2055116919856103.

79. Chen, J., Lee, K.H., Steinhauer, D.A., Stevens, D.J., Skehel, J.J., and Wiley, D.C. (1998). Structure of the hemagglutinin precursor cleavage site, a determinant of influenza pathogenicity and the origin of the labile conformation. Cell 95, 409–417.

80. Cheng, J., Zhao, Y., Xu, G., Zhang, K., Jia, W., Sun, Y., Zhao, J., Xue, J., Hu, Y., and Zhang, G. (2019). The S2 subunit of QX-type infectious bronchitis coronavirus spike protein is an essential determinant of neurotropism. Viruses 11, 972.

81. Claas, E.C., Osterhaus, A.D., van Beek, R., De Jong, J.C., Rimmelzwaan, G.F., Senne, D.A., Krauss, S., Shortridge, K.F., and Webster, R.G. (1998). Human influenza A H5N1 virus related to a highly pathogenic avian influenza virus. Lancet 351, 472–477.

82. Frana, M.F., Behnke, J.N., Sturman, L.S., and Holmes, K.V. (1985). Proteolytic cleavage of the E2 glycoprotein of murine coronavirus: host-dependent differences in proteolytic cleavage and cell fusion. J Virol 56, 912–920.

83. Kido, H., Okumura, Y., Takahashi, E., Pan, H.Y., Wang, S., Yao, D., Yao, M., Chida, J., and Yano, M. (2012). Role of host cellular proteases in the pathogenesis of influenza and influenza-induced multiple organ failure. Biochim Biophys Acta 1824, 186–194.

84. Le Coupanec, A., Desforges, M., Meessen-Pinard, M., Dube, M., Day, R., Seidah, N.G., and Talbot, P.J. (2015). Cleavage of a neuroinvasive human respiratory virus spike glycoprotein by proprotein convertases modulates neurovirulence and virus spread within the central nervous system. PLoS Pathog 11, e1005261.

85. Licitra, B.N., Millet, J.K., Regan, A.D., Hamilton, B.S., Rinaldi, V.D., Duhamel, G.E., and Whittaker, G.R. (2013). Mutation in spike protein cleavage site and pathogenesis of feline coronavirus. Emerg Infect Dis 19, 1066–1073.

86. Sun, X., Tse, L.V., Ferguson, A.D., and Whittaker, G.R. (2010). Modifications to the hemagglutinin cleavage site control the virulence of a neurotropic H1N1 influenza virus. J Virol 84, 8683–8690.

87. Benton, D.J., Wrobel, A.G., Xu, P., Roustan, C., Martin, S.R., Rosenthal, P.B., Skehel, J.J., Gamblin, S.J. (2020). Receptor binding and priming of the spike protein of SARS-CoV-2 for membrane fusion. Nature (2020). https://doi.org/10.1038/s41586-020-2772-0.

88. Li, Q., Wu, J., Nie, J., Zhang, L., Hao, H., Liu, S., Zhao, C., Zhang, Q., Liu, H., Nie, L., et al. (2020). The impact of mutations in SARS-CoV-2 spike on viral Infectivity and antigenicity. Cell 182, 1284–1294 e1289.

89. Phillips, J.M., Gallagher, T., and Weiss, S.R. (2017). Neurovirulent murine coronavirus JHM.SD uses cellular zinc metalloproteases for virus entry and cell-cell fusion. J Virol 91.

90. Giacca, M., Bussani, R., Schneider, E., Zentilin, L., Collesi, C., Ali, H., Braga, L., Secco, I., Volpe, M.C., Colliva, A., et al. (2020). Persistence of viral RNA, widespread thrombosis and abnormal cellular syncytia are hallmarks of COVID-19 lung pathology. medRxiv. doi: 10.1101/2020.06.22.20136358.

91. Greenlee, K.J., Werb, Z., and Kheradmand, F. (2007). Matrix metalloproteinases in lung: multiple, multifarious, and multifaceted. Physiol Rev 87, 69–98.

92. Grieve, A.G., and Rabouille, C. (2011). Golgi bypass: skirting around the heart of classical secretion. Cold Spring Harb Perspect Biol 3, a005298.

93. Harrison, S.C. (2008). Viral membrane fusion. Nat Struct Mol Biol 15, 690–698.

94. Roney, I.J., Rudner, A.D., Couture, J.F., and Kaern, M. (2016). Improvement of the reverse tetracycline transactivator by single amino acid substitutions that reduce leaky target gene expression to undetectable levels. Sci Rep 6, 27697.

95. Cockrell, A.S., and Kafri, T. (2007). Gene delivery by lentivirus vectors. Mol Biotechnol 36, 184–204.

